# Coupling Between Functionality and Trafficking to the Axon Initial Segment in KCNQ2/3 K^+^ Channels

**DOI:** 10.1101/2024.10.17.617761

**Authors:** Daisuke Yoshioka, Yasushi Okamura

**Affiliations:** Department of Physiology, Graduate School of Medicine, Osaka University, Suita, Osaka 565-0871, Japan

**Keywords:** KCNQ2/3, Ankyrin-G, Axon initial segment, Epilepsy, Lateral diffusion, Exocytosis, Endocytosis, Single-molecule imaging

## Abstract

KCNQ2/3 are the predominant voltage-gated K^+^ channels localized at the axon initial segment (AIS), the critical site for the initiation of action potentials and the plasticity of excitability. Both the functionality and spatial distribution of KCNQ2/3 provide the basis for the regulation of neuronal excitability and the pathogenesis of various neurological disorders, including epilepsy. However, less is known about how functionality is coupled with trafficking regulation in KCNQ2/3. Here, we study the AIS localization of KCNQ2/3 by performing both multiple- and single-molecule imaging analyses. We found that low-activity mutations in the KCNQ3 subunit affect all of the 3D dynamics composed of lateral diffusion and exo/endocytosis processes through disruption of interaction with ankyrin-G, consequently suppressing the AIS targeting of KCNQ2/3. Thus, the functionality of KCNQ2/3 is coupled with its trafficking regulation, enhancing our understanding of the mechanisms underlying physiological and pathophysiological changes in neuronal excitability.

## Introduction

The KCNQ2/3 is a type of neuronal voltage-gated potassium (Kv) channel. The KCNQ2/3-current is integral to the regulation and maintenance of membrane potential and firing rates of action potentials in neurons (Brown and Adams 1980; Wang et al. 1998; Shah et al. 2008). The role of KCNQ2/3-current in neuronal excitability is also pathologically significant because dysfunction of the KCNQ2/3 precipitates various neurological disorders, such as benign familial neonatal convulsions (BFNC) and early infantile epileptic encephalopathy (EIEE) (Biervert et al. 1998; Orhan et al. 2014; Nappi et al. 2020). Therefore, the KCNQ2/3 has been studied as an important therapeutic target for the neurological disorders (X. Li et al. 2021; T. Li et al. 2021).

Impact of KCNQ2/3-current on shaping patterns of neuronal firings is mediated through, not only the channel gating, but also the channel localization at specific subcellular sites such as the axon initial segment (AIS) and node of Ranvier (Pan et al. 2006; Rasmussen et al. 2007). Importantly, the spatial pattern and ion channel composition at the AIS play critical roles in the neuronal excitability. The AIS also shows morphological and functional plasticity; it changes flexibly in response to the neuronal activity (Grubb and Burrone 2010; Kuba, Oichi, and Ohmori 2010). Pathologically, defects of AIS structure and composition cause various types of neurological and psychiatric diseases such as epilepsy, neurodegeneration, bipolar disorder, and schizophrenia (Wimmer et al. 2010; Buffington and Rasband 2011; Hsu, Nilsson, and Laezza 2014).

In general, the spatial distribution of membrane proteins on the cell surface is generated and maintained via several trafficking pathways comprising lateral diffusion and exo/endocytosis. All the pathways potentially contribute to the spatial pattern formation of AIS proteins. In fact, there are several reports of different trafficking regulation mechanisms: 1) Nav1.6 channels being directly inserted into the AIS (Akin et al. 2015); 2) the concentration of Nav1.2 channels at the AIS being due to selective endocytosis at the somatodendritic regions (Fache et al. 2004); and 3) AIS proteins such as KCNQ3 and neurofascin186 being inserted into the soma and axon terminal and then reaching the AIS by lateral diffusion on the cell surface in immature neurons (Ghosh et al. 2020). However, it remains unclear what kinds of trafficking mechanisms regulate the spatial distribution of KCNQ2/3 in mature neurons.

In KCNQ2/3, many intramolecular modules have been suggested to be involved in the regulation of functionality and trafficking. Regarding KCNQ2/3 functionality, recent studies on the protein structure of KCNQ2 have led to a detailed understanding of the functional mechanisms of KCNQ2/3 at the atomic level (X. Li et al. 2021; Ma et al. 2023; Zhang et al. 2024). Specifically, S1-S4 helices form the voltage-sensing domain (VSD), and the S4-S5 linker and the edge of S5-S6 helices contain multiple protein-protein interaction sites and PI(4,5)P_2_ binding sites which are necessary for electro-mechanical (E-M) coupling of KCNQ2/3 (Yang et al. 2022; Okamura and Yoshioka 2023). In contrast, regarding the KCNQ2/3 trafficking, it is critical to interact with ankyrin-G, which is a major scaffold protein underlying the AIS, via the ankyrin binding motif (ABM) at the end of the C-terminal cytoplasmic region of KCNQ2/3 (Pan et al. 2006; Rasmussen et al. 2007). The other trafficking regulatory modules mainly overlap with the activity regulatory modules, such as the pore domain (PD) and the C-terminal cytoplasmic region including A-D helices which are responsible for PI(4,5)P_2_ binding, CaM interaction, and tetramer formation of KCNQ2/3 (Chung, Jan, and Jan 2006; Haitin and Attali 2008; Abidi et al. 2015; Kim et al. 2018; Hefting et al. 2020). These findings raise the possibility that KCNQ2/3 functionality and trafficking are not completely independent of each other. However, the interplay between the channel functionality and trafficking has not been fully studied and it still remains elusive how the functionality of KCNQ2/3 is coupled with its trafficking process in neurons.

Here, we analyzed the trafficking process of KCNQ2/3 in living neurons using multiple- and single-molecule imaging. Results of the three-dimensional dynamics of KCNQ2/3 provided evidence that the interaction with ankyrin-G affects both lateral diffusion and exo/endocytosis of KCNQ2/3 at the AIS regions in neurons. Furthermore, we found that several mutations of sites important for KCNQ2/3 functionality also altered all the three-dimensional dynamics, consequently inducing less AIS localization of KCNQ2/3. The altered spatial dynamics of the low-activity KCNQ2/3 were suggested to be associated with weakened interaction with ankyrin-G. In conclusion, this study revealed the coupling between functionality and trafficking regulation of KCNQ2/3 in neurons.

## Results

### Halo-TAC-KCNQ3 enables verification of the functionality and trafficking of KCNQ2/3

To directly measure the spatial dynamics of KCNQ2/3 in living neurons, we generated a mouse KCNQ3 fused with a HaloTag (Figure 1A), since KCNQ3, but not KCNQ2, has been shown to be a predominant trafficking regulator of KCNQ2/3 (Rasmussen et al. 2007). According to a previous report (Benned-Jensen et al. 2016), we designed the construct in which HaloTag is expressed in the extracellular region by adding a single-pass transmembrane domain derived from IL2RA (TAC) to the N-terminus of KCNQ3 (Halo-TAC-KCNQ3).

**Figure 1.**
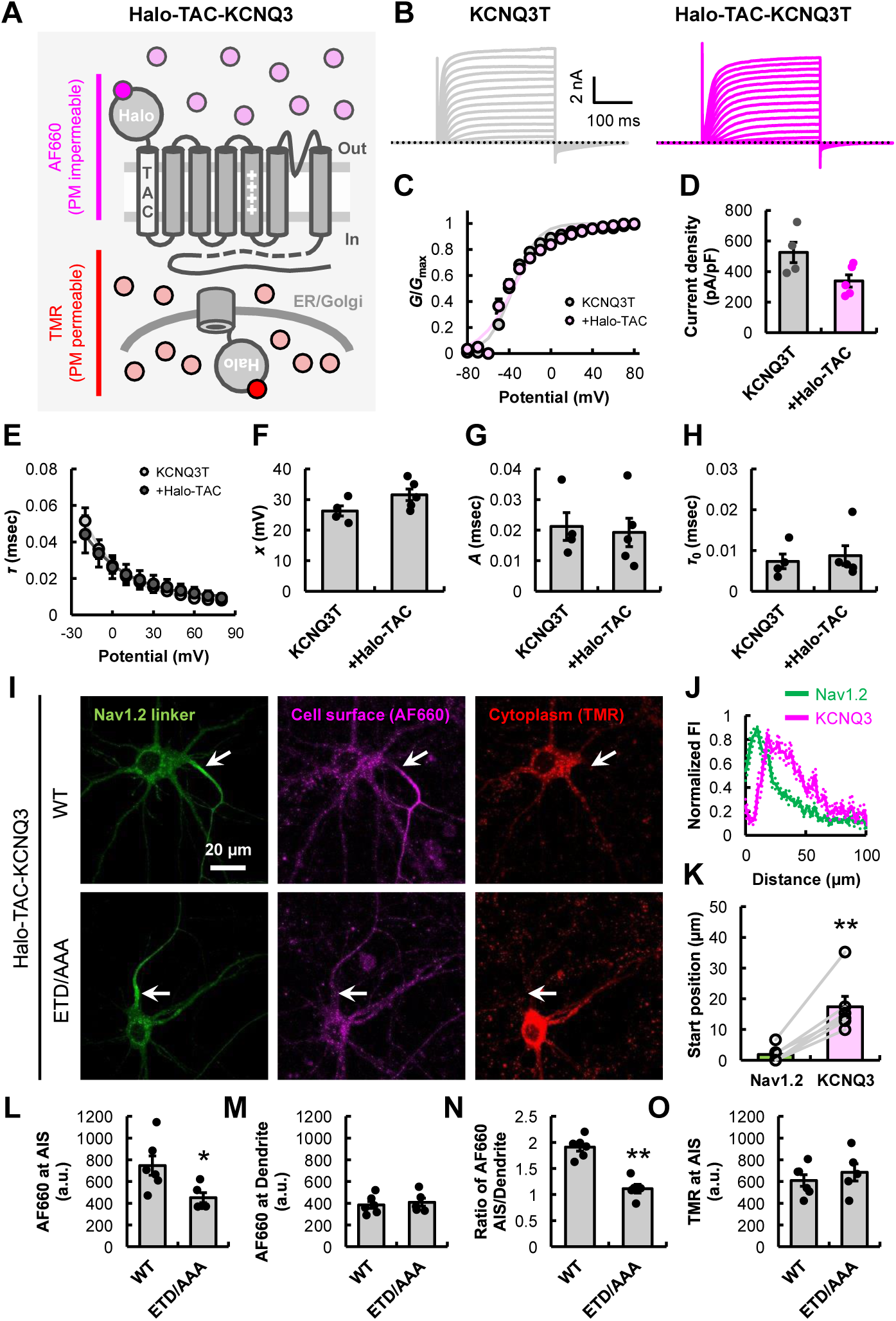
Halo-TAC-KCNQ3 enables verification of the functionality and trafficking of KCNQ2/3. (A) Schematic representation of Halo-TAC-KCNQ3. (B) Representative current traces of KCNQ3T or Halo-TAC-KCNQ3T obtained by whole-cell patch-clamp recordings in HEK293T cells. (C) *G*-*V* curves of KCNQ3T and Halo-TAC-KCNQ3T. *V*_1/2_ for KCNQ3T: -35.11 ± 1.67 mV (*n* = 4) and for Halo-TAC-KCNQ3T: -35.38 ± 1.87 mV (*n* = 5). *P* = 0.927 (Welch’s *t*-test). (D) Current densities at 80 mV for KCNQ3T (*n* = 4) and Halo-TAC-KCNQ3T (*n* = 5). *P* = 0.090 (Welch’s *t*-test). (E) Activation time constant (*τ*) of KCNQ3T (*n* = 4) and Halo-TAC-KCNQ3T (*n* = 5) at different potentials. (F-H) The quantified parameters: *x* (F, *P* = 0.105), *A* (G, *P* = 0.801), and *τ*_0_ (H, *P* = 0.692). *P*-values were obtained by Welch’s *t*-test. (I) Representative confocal microscopic images of primary cultured hippocampal neurons expressing EGFP-Nav1.2 II-III linker and Halo-TAC-KCNQ3 with KCNQ2. Halo-TAC-KCNQ3 (WT, ETD/AAA) was labeled with AF660 and TMR. The arrowhead indicates the position of AIS. Scale bar, 20 μm. (J and K) Average intensity profiles of EGFP-Nav1.2 II-III linker and Halo-TAC-KCNQ3 (J), and their start positions from the edge of the soma (K, \*\**P* = 0.002 by paired *t*-test) in neurons (*n* = 6). (L-O) Average AF660 intensity at the AIS (L, \**P* = 0.029) and dendrite (M, *P* = 0.266), ratio of AF660 intensity between AIS and dendrite (N, \*\**P* = 0.000), TMR intensity at the AIS (O, *P* = 0.501) of neurons expressing Halo-TAC-KCNQ3 (WT, *n* = 6; ETD/AAA, *n* = 5). *P*-values were obtained by Welch’s *t*-test. Data are mean ± SE.

First, we checked the channel functionality of Halo-TAC-KCNQ3 by heterologous expression in HEK293T cells. Only in electrophysiological experiments to verify the properties of the homomeric KCNQ3 channels, we utilized a KCNQ3-A316T mutant (KCNQ3T) that allows KCNQ3 to conduct current as a homotetramer (Etxeberria et al. 2004). The currents of KCNQ3T or Halo-TAC-KCNQ3T were recorded by whole-cell patch clamp in HEK293T cells. The results indicate that the channel properties such as current density, *V*_1/2_, and activation kinetics of Halo-TAC-KCNQ3T were similar to those of KCNQ3T (Figures 1B-1H).

Next, we verified the subcellular spatial distribution of Halo-TAC-KCNQ3 in neurons. Halo-TAC-KCNQ3 was co-expressed with mouse KCNQ2 in cultured hippocampal neurons at 10 days *in vitro* (DIV) which represent mature neurons (JoséGarrido et al. 2003; Bréchet et al. 2008). Then, Halo-TAC-KCNQ3 was sequentially labeled with cell membrane-impermeable and - permeable HaloTag ligands (AF660 and TMR), in order to differentially detect KCNQ2/3 present on the cell surface and in the cytoplasmic pool at the same time. The AIS region was identified by heterologous expression of EGFP-Nav1.2 II-III linker as the AIS marker (JoséGarrido et al. 2003; Bréchet et al. 2008). The fluorescence signal showed that Halo-TAC-KCNQ3 was uniformly distributed throughout the cytoplasm, while on the cell surface, it was predominantly localized at the AIS region (Figure 1I). In particular, as previously reported for endogenous KCNQ3 (Pan et al. 2006; Van Wart, Trimmer, and Matthews 2007; Yamada and Kuba 2016), the start position of the Halo-TAC-KCNQ3 distribution was shifted to the distal side of the AIS region from the edge of the soma (17.40 ± 3.38 μm) compared to the AIS marker (1.86 ± 0.97 μm) (Figures 1J and 1K).

Additionally, we investigated whether the trafficking of Halo-TAC-KCNQ3 is regulated by its interaction with ankyrin-G. To this end, we examined the spatial pattern of KCNQ3-E838A/T839A/D840A (ETD/AAA), a mutant in which the amino acid residues forming ABM are altered (Pan et al. 2006; Rasmussen et al. 2007). As anticipated, the KCNQ3-ETD/AAA mutant was not efficiently expressed on the cell surface specifically at the AIS (Figures 1I and 1L-1N), despite being well expressed in the cytoplasmic region similar to the wild-type (Figure 1O). This suggests that the trafficking of Halo-TAC-KCNQ3 is regulated via interactions with ankyrin-G, as endogenous KCNQ3 (Pan et al. 2006; Rasmussen et al. 2007). These results suggest that the expression of Halo-TAC-KCNQ3 recapitulates that of endogenous KCNQ3 in cultured hippocampal neurons, thereby establishing the validity of Halo-TAC-KCNQ3 for analyzing the functionality and trafficking of KCNQ2/3 in neurons.

In addition, we also constructed Halo-TAC-KCNQ2. We verified that Halo-TAC-KCNQ2 has similar channel properties to KCNQ2 (Figures S1A-S1G). As in Halo-TAC-KCNQ3, Halo-TAC-KCNQ2 was normally localized at the cell surface of the AIS (Figures S1H and S1J-S1L). Co-expression of KCNQ3-ETD/AAA inhibited the AIS targeting of Halo-TAC-KCNQ2 more strongly than mutation of the ABM of Halo-TAC-KCNQ2 (Figures S1H-S1L). These results support the idea that KCNQ3 is a dominant trafficking regulator of KCNQ2/3 (Rasmussen et al. 2007), highlighting the reason why we focused on KCNQ3 to analyze the spatial dynamics of KCNQ2/3.

### Targeting efficiency of KCNQ2/3 at the AIS correlates with their channel functionality

To investigate the relationships between functionality and spatial dynamics of KCNQ2/3 using the constructed Halo-TAC-KCNQ3, several mutations were introduced into the activity regulatory site of KCNQ3 to manipulate channel functionality. For the manipulation of VSD activation, we mutated the triple amino acid residues that constitute the VSD, namely KCNQ3T-F168R/D203N/Q234E (KCNQ3T-FDQ/RNE) to inhibit the transition of VSD conformation from the resting state to the activated state (Yang et al. 2022). The KCNQ3T-FDQ/RNE mutant was not activated and exhibited negligible current (Figures 2A and 2B). For the manipulation of E-M coupling, several mutations (W248R, Y267C, and S343F in mouse) were introduced to KCNQ3 (Yang et al. 2022). Notably, the Y267C mutation of mouse KCNQ3, corresponding to the Y266C mutation of human KCNQ3, is also associated with EIEE (Symonds et al. 2019) (ClinVar accession ID: VCV000205964.2). Indeed, these mutants also exhibited negligible current (Figures 2A and 2B). Additionally, it is known that PI(4,5)P_2_, a type of inositol phospholipid, directly controls the functionality of various ion channels including KCNQ2/3 (Hilgemann and Ball 1996; Delmas and Brown 2005; Suh and Hille 2008; Okamura and Yoshioka 2023). We mutated basic residues constituting the PI(4,5)P_2_ binding site belonging to the S4-S5 linker and the end of S5 and S6 helices of KCNQ3, namely KCNQ3T-R243A, KCNQ3T-H258N, KCNQ3T-K260A, and KCNQ3T-K359A/R365A/K367A (KCNQ3T-KRK/AAA) (Zhou et al. 2013; Choveau et al. 2018). These mutations of PI(4,5)P_2_ binding sites significantly decreased the current density and shifted *V*_1/2_ of KCNQ3T to a more positive range (Figures 2A-2D). In particular, the mutation of the H258 residue, corresponding to H257 in human KCNQ3, drastically reduces the current density of KCNQ3T, and the H257Y mutation of human KCNQ3 is also associated with EIEE (ClinVar accession ID: VCV000937869.5).

**Figure 2.**
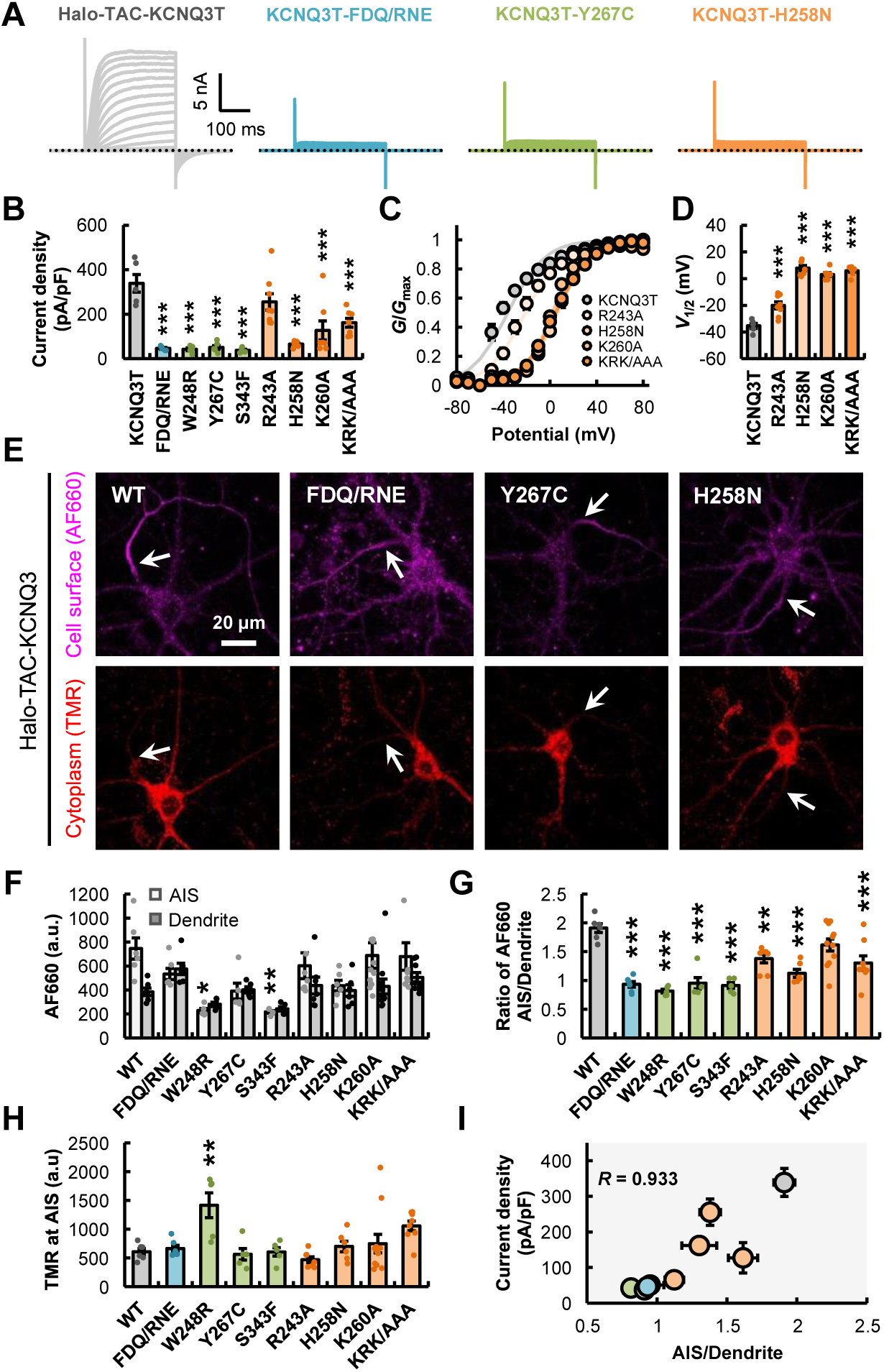
Targeting efficiency of KCNQ2/3 at the AIS correlates with their channel functionality. (A) Representative current traces of Halo-TAC-KCNQ3T or the indicated mutants obtained by whole-cell patch-clamp recordings in HEK293T cells. (B-D) Current densities at 80 mV (B), *G*-*V* curves (C) and quantified *V*_1/2_ (D) for Halo-TAC-KCNQ3T or the mutants (*n* = 5-8). \*\**P* < 0.01 and \*\*\**P* < 0.001 (Dunnett’s test, comparisons were made with the “KCNQ3T” column). (E) Representative confocal microscopic images of primary cultured hippocampal neurons expressing Halo-TAC-KCNQ3 or the mutants. Halo-TAC-KCNQ3 was labeled with AF660 and TMR. The arrowhead indicates the position of AIS. Scale bar, 20 μm. (F-H) Average AF660 intensity at the AIS and dendrite (F), ratio of AF660 intensity between AIS and dendrite (G), TMR intensity at the AIS (H) of neurons expressing WT or the mutant Halo-TAC-KCNQ3 (*n* = 5-11). \**P* < 0.05, \*\**P* < 0.01, and \*\*\**P* < 0.001 (Dunnett’s test, comparisons were made with the “WT” column for the same group). (I) Correlation between current density (B) and ratio of AF660 intensity between AIS and dendrite (G). Spearman’s rank correlation coefficient (*R*) = 0.933. Data are mean ± SE.

Next, we examined the effect of these mutations leading to the alteration of functionality of KCNQ3 on the trafficking in neurons. We co-expressed wild-type or mutant Halo-TAC-KCNQ3 with wild-type KCNQ2 in cultured hippocampal neurons and performed confocal microscopy imaging (Figure 2E). To quantitatively compare these spatial patterns between wild-type and mutants, we analyzed the AF660 signal indicating the surface density of Halo-TAC-KCNQ3 in different intracellular regions such as AIS and dendrites. Interestingly, it was found that the surface density of low-activity KCNQ3 mutants was significantly suppressed at the AIS region but not the dendrites (Figures 2F and 2G), although the protein expression level of all mutants was not decreased in the cytoplasm (Figure 2H). Surprisingly, the AIS selectivity, which was defined as the ratio of fluorescence intensity at the dendrite to AIS, obviously correlated with the current density of KCNQ3 measured in HEK293T cells (Figure 2I). In addition, the expression of low-activity mutant KCNQ3 (KCNQ3-H258N) also suppressed the AIS targeting of Halo-TAC-KCNQ2 in a dominant-negative manner (Figures S1H and S1J-S1L). These results provide evidence that the trafficking dynamics of KCNQ2/3 are regulated in a channel functionality-dependent manner.

### Heteromer formation with KCNQ2 is not prevented by the low-activity mutations of KCNQ3

To understand the molecular mechanisms underlying the coupling between functionality and trafficking regulation of KCNQ2/3, we examined the heteromer formation of KCNQ2/3 because KCNQ2 and KCNQ3 homomers are not normally transported to the cell surface in neurons (Figures 3A-3C, S1H, and S1J-S1L) (Rasmussen et al. 2007). These data support that, under Halo-TAC-KCNQ3 and KCNQ2 co-expression, the fluorescent signal detected on the cell surface primarily reflects heteromeric KCNQ2/3 complexes, but not KCNQ3 homomers. In addition, these data also propose the possibility that mutations in KCNQ3 unexpectedly inhibit KCNQ2/3 heteromer formation, resulting in reduced AIS selectivity.

**Figure 3.**
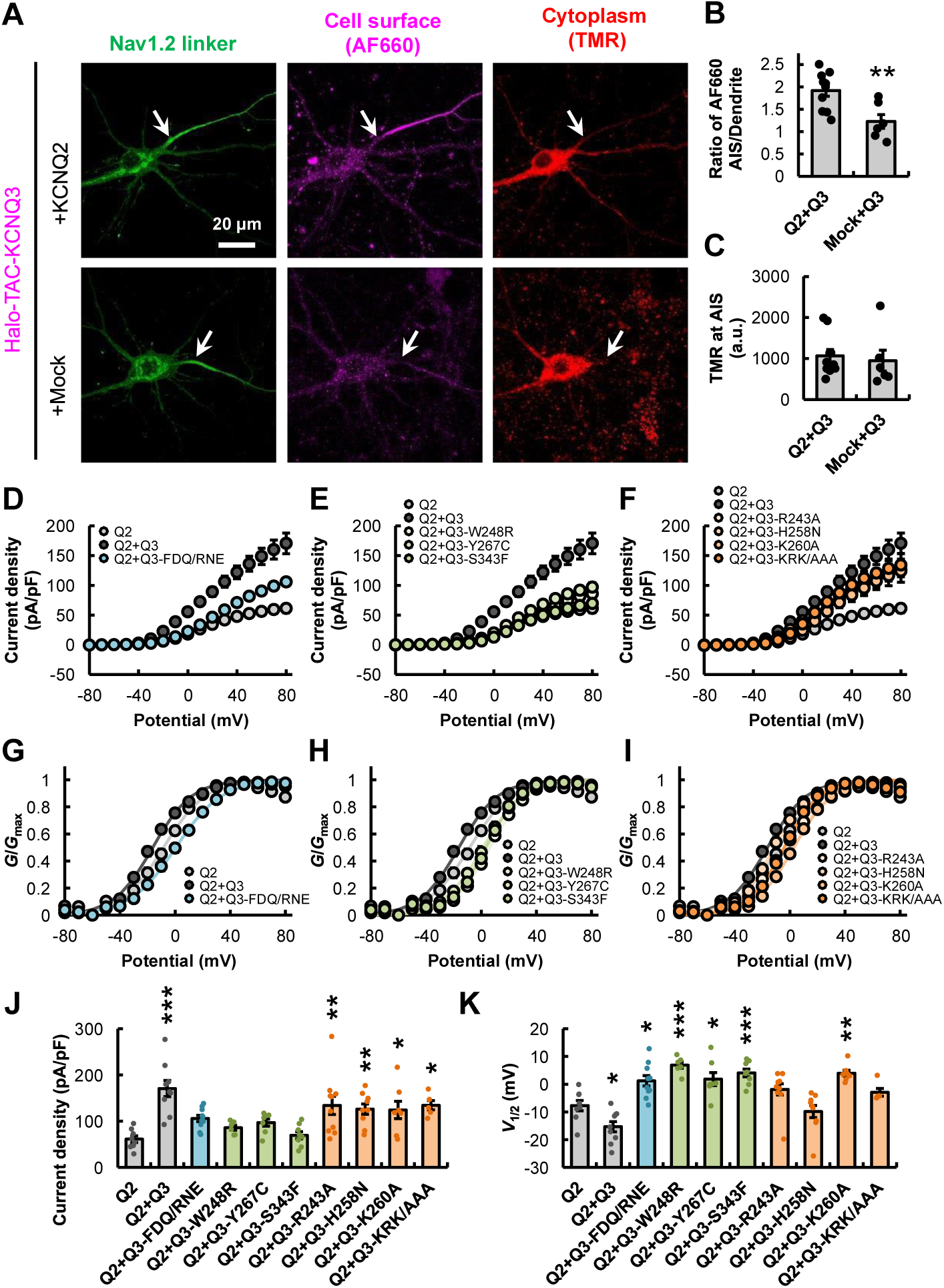
Heteromer formation with KCNQ2 is not prevented by the low-activity mutations of KCNQ3. (A) Representative confocal microscopic images of primary cultured hippocampal neurons expressing EGFP-Nav1.2 II-III linker and Halo-TAC-KCNQ3 with KCNQ2 or empty vector. Halo-TAC-KCNQ3 was labeled with AF660 and TMR. Scale bar, 20 μm. (B and C) Ratio of AF660 intensity between AIS and dendrite (B, \*\**P* = 0.008), TMR intensity at the AIS (C, *P* = 0.722) of neurons expressing Halo-TAC-KCNQ3 with KCNQ2 (*n* = 10) or empty vector (*n* = 6). *P*-values were obtained by Welch’s *t*-test. (D-K) I-V curves (D-F) and G-V curves (G-I) derived from whole-cell patch-clamp recordings in HEK293T cells expressing either KCNQ2 or a heteromeric complex of KCNQ2 and Halo-TAC-KCNQ3. The current density at 80 mV (J) and *V*_1/2_ (K) were quantified (*n* = 5-10). \**P* < 0.05, \*\**P* < 0.01, and \*\*\**P* < 0.001 (Dunnett’s test, comparisons were made with the “Q2” column). Data are mean ± SE.

To test this possibility, we co-expressed wild-type or mutant Halo-TAC-KCNQ3 with wild-type KCNQ2 in HEK293T cells and performed whole-cell patch clamp recording. KCNQ2/3 heteromers showed a larger current than KCNQ2 homomers. Importantly, four KCNQ3 mutants including H258N also showed a significant current increase when co-expressed with KCNQ2 compared to KCNQ2 homomers (Figures 3D-3F and 3J). The expression of other KCNQ3 mutants also significantly shifted the *V*_1/2_ of KCNQ2/3-current to a positive range compared to wild-type KCNQ2 homomers and KCNQ2/3 heteromers (Figures 3G-3I and 3K). These results suggest that low-activity KCNQ3 mutants can still form the heteromeric KCNQ2/3 complex. Therefore the decrease in AIS selectivity of low-activity KCNQ3 mutants is not due to a disruption of heteromeric KCNQ2/3 complex formation.

### Neuronal excitability and AIS structure are not affected by the expression of low-activity KCNQ3 mutants

Next, we measured the intrinsic excitability of neurons co-expressing wild-type or H258N mutant Halo-TAC-KCNQ3 with KCNQ2, because neuronal excitability is critical for the spatial pattern of ion channels at the AIS (Grubb and Burrone 2010; Kuba, Oichi, and Ohmori 2010; Lezmy et al. 2017). The results of whole-cell current clamp recordings showed that there was no significant difference in the average firing rate and the resting membrane potential (RMP) between wild-type and H258N mutant (Figures 4A-4C). In addition, we also analyzed the waveform of action potentials and found no significant difference in both amplitude and half-width of action potentials (Figures 4D-4G).

**Figure 4.**
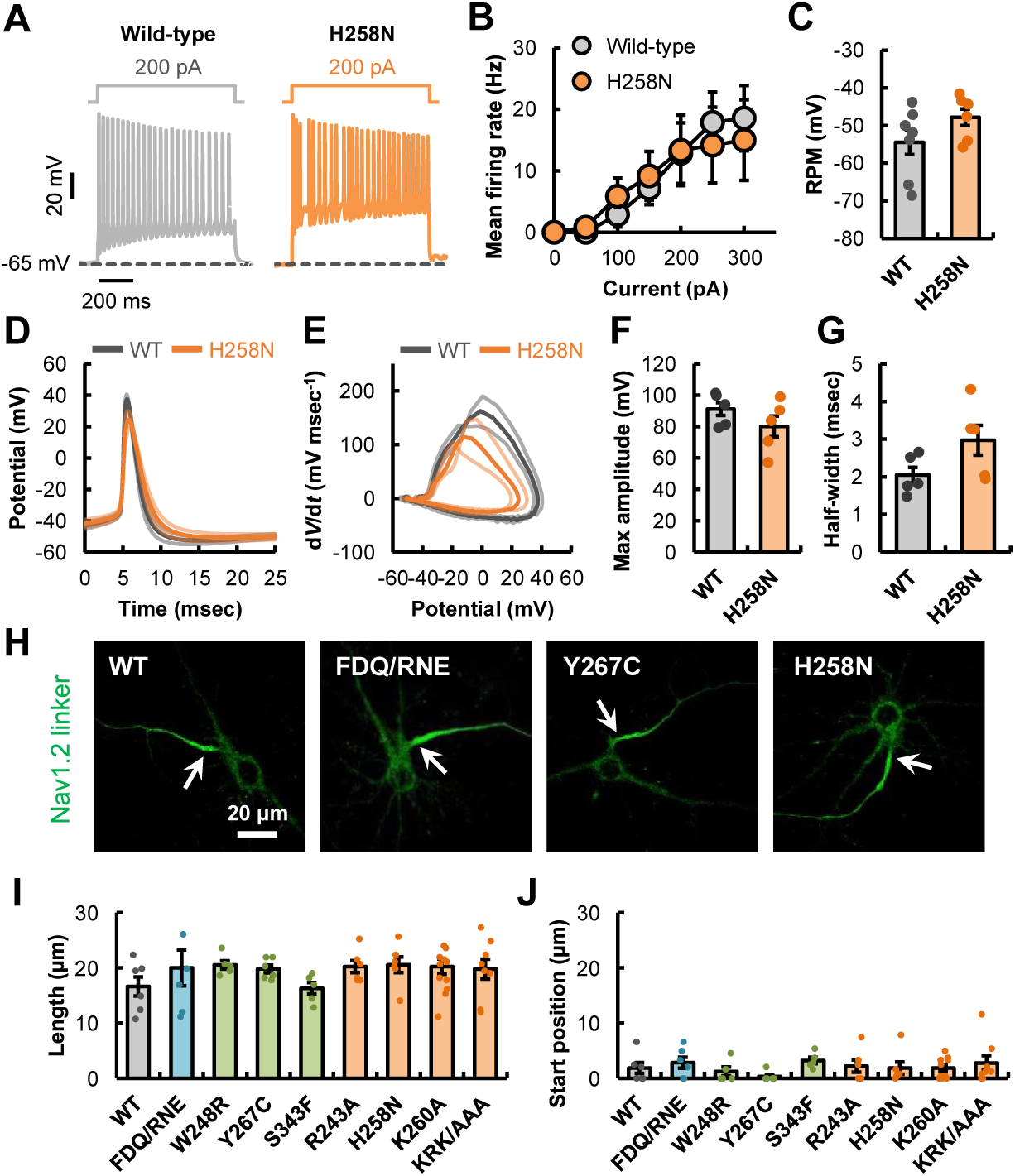
Neuronal excitability and AIS structure are not affected by the expression of low-activity KCNQ3 mutants. (A) Whole-cell patch clamp recording from primary cultured hippocampal neurons expressing Halo-TAC-KCNQ3 or Halo-TAC-KCNQ3-H258N with KCNQ2. Representative action potentials evoked by 200 pA current injection for 800 msec. (B) Mean firing rate evoked by current injection with increments of 50 pA in neurons expressing Halo-TAC-KCNQ3 (*n* = 7) or Halo-TAC-KCNQ3-H258N (*n* = 6) (50 pA, *P* = 0.363; 100 pA, *P* = 0.466; 150 pA, *P* = 0.707; 200 pA, *P* = 0.599; 250 pA, *P* = 0.677; 300 pA, *P* = 0.706). *P*-values were obtained by Welch’s *t*-test. (C) Resting membrane potential (RMP). *P* = 0.151 (Welch’s *t*-test). (D-G) Average waveform (D), phase plot (E), maximum amplitude (F, *P* = 0.249), and half-width (G, *P* = 0.114) of spontaneous action potentials at resting membrane potential in neurons expressing Halo-TAC-KCNQ3 (*n* = 5) or Halo-TAC-KCNQ3-H258N (*n* = 5). *P*-values were obtained by Welch’s *t*-test. (H) Representative confocal microscopic images of primary cultured hippocampal neurons expressing EGFP-Nav1.2 II-III linker with KCNQ2 and Halo-TAC-KCNQ3 (WT or the mutants). The arrowhead indicates the position of AIS. Scale bar, 20 μm. (I) The lengths of EGFP-Nav1.2 II-III linker localized region (*n* = 5-12). *P* > 0.05 for wild-type versus all mutant KCNQ3 (Dunnett’s test, comparisons were made with the “WT” column). (J) The start positions of EGFP-Nav1.2 II-III linker localized region from the edge of the soma in neurons (*n* = 5-12). *P* > 0.05 for wild-type versus all mutant KCNQ3 (Dunnett’s test, comparisons were made with the “WT” column). Data are mean ± SE.

We also verified the effect of mutant KCNQ3 expression on the AIS structure defined by the AIS marker (EGFP-Nav1.2 II-III linker). In this experiment, wild-type or mutant Halo-TAC-KCNQ3, KCNQ2, and the AIS marker were expressed in hippocampal neurons (10 DIV). Based on the signal of the AIS marker, the morphological properties such as length and start position of AIS structure were quantified respectively. As a result, it was confirmed that the spatial pattern of AIS markers was not rearranged by the expression of KCNQ3 mutants (Figures 4H-4J). These results indicate that heterologous expression of mutant KCNQ3 does not affect the neuronal excitability and the AIS structure in our experimental setup.

### Lateral diffusivity of KCNQ2/3 at the AIS is enhanced by the low-activity mutation of KCNQ3

Generally, the spatial pattern of membrane proteins is formed by the three-dimensional trafficking pathway composed of exocytosis, lateral diffusion, and endocytosis. Therefore, the change in AIS localization observed in low-activity KCNQ3 mutants is presumed to be the result of a change in any of these three types of trafficking pathways. To analyze the detailed trafficking process of KCNQ2/3 with high spatiotemporal resolution, we performed single-molecule imaging of TMR-labeled Halo-TAC-KCNQ3 with Total Internal Reflection Fluorescence Microscopy (TIRFM) (M J Saxton and Jacobson 1997; Axelrod 2003), and succeeded in detecting Halo-TAC-KCNQ3 with high specificity at the single-molecule level (Figures 5A and S2A-S2F; Videos S1-S3).

**Figure 5.**
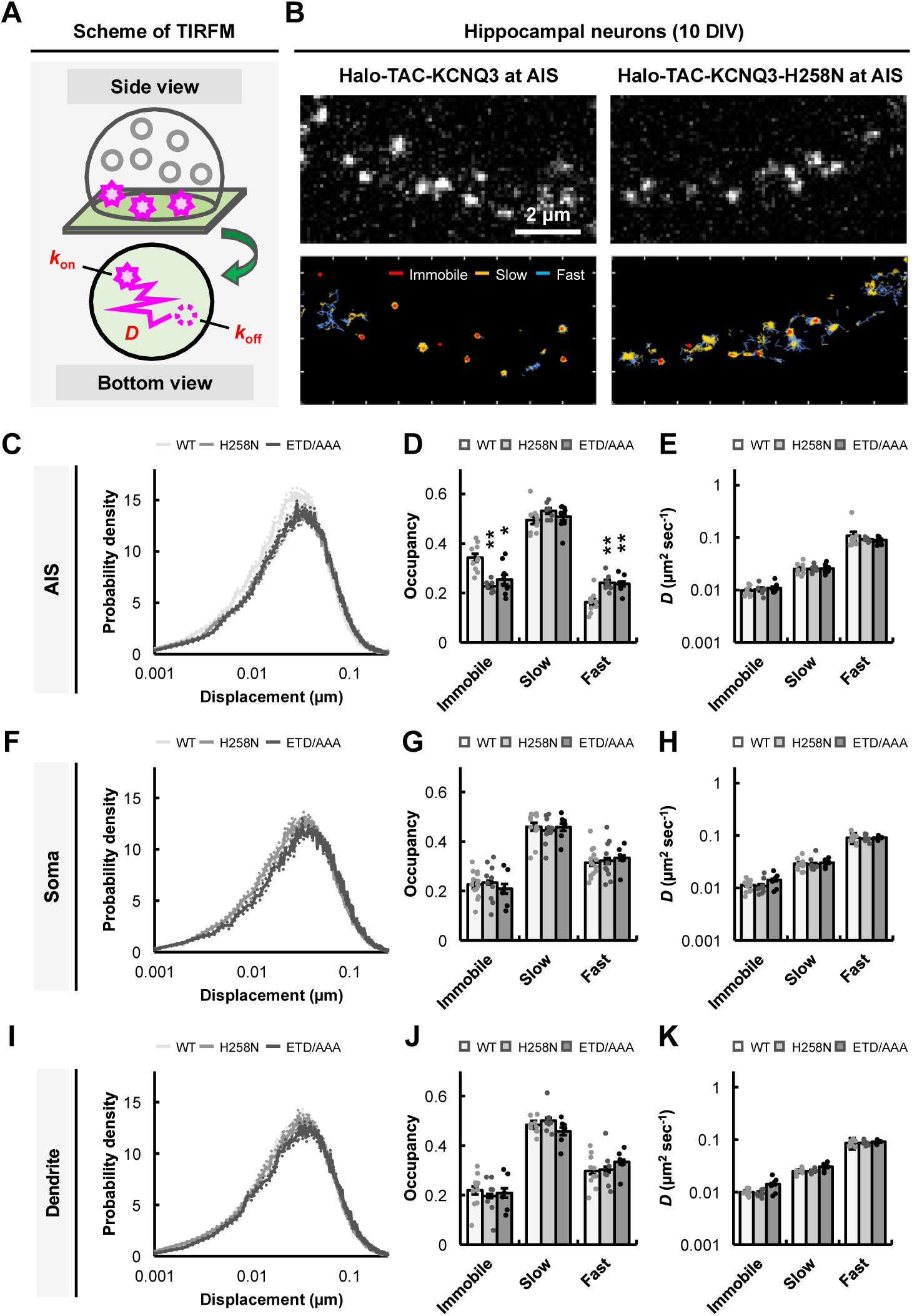
Lateral diffusivity of KCNQ2/3 at the AIS is enhanced by the low-activity mutation of KCNQ3. (A) A schematic diagram illustrating single-molecule imaging using TIRFM. (B) Representative TIRFM images (top) and the diffusion trajectories for 40 sec (bottom) of primary cultured hippocampal neurons expressing Halo-TAC-KCNQ3 (WT, H258N) labeled with TMR at the AIS. Trajectories of 3 diffusion states of the molecule within a unit time interval of 33 msec are shown by different colors. Scale bar, 2 μm. (C-K) The probability density distributions of the displacement during 33 msec (C, F, and I), the occupancies (D, G, and J), and the diffusion coefficients (E, H, and K) of Halo-TAC-KCNQ3 at the AIS (WT, *n* = 10; H258N, *n* = 7; ETD/AAA, *n* = 9), the soma (WT, *n* = 14; H258N, *n* = 10; ETD/AAA, *n* = 6), and the dendrite (WT, *n* = 10; H258N, *n* = 7; ETD/AAA, *n* = 6). \**P* < 0.05 and \*\**P* < 0.01 (Steel-Dwass test, only comparisons with the “WT” group for the same state are depicted). Data are mean ± SE.

At first, we analyzed the lateral diffusion dynamics from the diffusion trajectories of the KCNQ2/3 (Figure 5B). A sum of two-dimensional diffusion equations was used to fit the displacement distribution of Halo-TAC-KCNQ3 (Figures 5C, 5F, and 5I). The optimal number of diffusion states was estimated to be three by the Akaike Information Criterion (AIC) (Figures S2G-S2I; see “Methods” for Analysis of single-molecule dynamics). Then, using a hidden Markov model (HMM), the diffusion trajectories of KCNQ2/3 were classified into 3 states, immobile, slow-diffusion, and fast-diffusion states, with different diffusion coefficients. The occupancy of the immobile state at the AIS (34%) was significantly higher than the one at the soma (23%) and dendrite (22%), whereas the occupancy of the fast-diffusion state at the AIS (16%) was significantly lower than the one at the soma (31%) and dendrite (30%) (Figures 5D, 5G, and 5J). The diffusion coefficients of wild-type Halo-TAC-KCNQ3 did not change between the AIS and somatodendritic regions (Figures 5E, 5H, and 5K). Furthermore, Mean Square Displacement (MSD) analysis showed that the confinement radius of the slow-diffusion state of KCNQ3 was extended from 70 nm at the AIS to 298 nm at the soma and 257 nm at the dendrite (Figure S3). These data indicate that the diffusion of KCNQ3 is more strongly confined at the AIS compared with the somatodendritic region.

We also analyzed two variants of KCNQ3: KCNQ3-H258N, a mutant of epilepsy-associated residues (ClinVar accession ID: VCV000937869.5) with apparently low AIS selectivity, and KCNQ3-ETD/AAA, which lacks ankyrin-G binding affinity. The immobile state occupancy of KCNQ3-H258N and KCNQ3-ETD/AAA at the AIS significantly decreased to 23% and 24% respectively, whereas the fast-diffusion state occupancy increased to 24% and 25% respectively, thus showing the same diffusion behavior as at the somatodendritic region (Figures 5D, 5G, and 5J). The diffusion coefficient and the confinement radius in each state of both mutants were equivalent to those of wild-type KCNQ3 in all subcellular regions (Figures 5E, 5H, 5K, and S3). The results of KCNQ3-H258N show that the activity regulatory site affects the diffusivity of KCNQ2/3 at the AIS. In addition, the results of KCNQ3-ETD/AAA indicate that the interaction between ankyrin-G and KCNQ2/3 is one of the factors determining the immobile and fast-diffusion states.

### Exo/endocytosis of KCNQ2/3 at the AIS is altered by the low-activity mutation of KCNQ3

In addition to the lateral diffusion, we also analyzed the exo/endocytosis process of KCNQ2/3 in neurons. The appearance and disappearance of bright spots obtained by single-molecule imaging indicate the insertion of molecules into the membrane by exocytosis and the internalization of molecules by endocytosis, respectively (Figure 6A). First, we calculated the exocytosis rate constant (*k*_on_) as the density of molecules newly appearing on the cell surface per unit time. The analysis revealed that the *k*_on_ of wild-type KCNQ3 increased at the AIS compared to the somatodendritic regions (Figures 6B and 6C). The H258N and ETD/AAA mutations decreased the *k*_on_ at the AIS, but not at the somatodendritic region (Figures 6B and 6C). Here, it should be noted that the *k*_on_ can be affected by the molecular concentration of KCNQ3 in the cytoplasmic pool. However, we have confirmed that the expression amounts of KCNQ3 in the cytoplasm were not affected by the H258N and ETD/AAA mutations (Figures 1O and 2H). This enables the comparison of the *k*_on_ of KCNQ3 between the wild-type and mutants (KCNQ3-H258N and KCNQ3-ETD/AAA), disregarding the effects of protein expression levels in the cytoplasm. Next, we quantified the endocytosis process. Since the self-transition probability of Halo-TAC-KCNQ3 was high (83∼98%), each molecule was classified into immobile, slow-, and fast-diffusing molecules based on the diffusion state with the highest occupancy in the trajectory of the molecule (Figure S4). Given the high self-transition probability of Halo-TAC-KCNQ3, the occupancy of each diffusion state (Figure 5) is mainly accounted for by the endocytosis rate constant (*k*_off_) and/or the fraction of each classified group. The *k*_off_ was calculated as the inverse of the average lifetime of KCNQ3 in each classified group. Interestingly, the *k*_off_ of KCNQ3 correlated with the diffusivity of each molecule in all subcellular regions: the slower the diffusion, the longer the lifetime on the cell surface (Figures 6D-6I). However, there is no significant difference in the *k*_off_ of KCNQ3 between the AIS and somatodendritic regions, and between the wild-type and mutants (Figures 6D-6I). In contrast, the fraction of fast-diffusing KCNQ3 with the shortest lifetime increased at the somatodendritic region compared with the AIS (Table S1). The H258N and ETD/AAA mutations decreased the fraction of immobile KCNQ3 with the longest lifetime and increased the fraction of fast-diffusing KCNQ3 with the shortest lifetime at the AIS (Table S1). Taken together, our data on KCNQ3-H258N indicate that the low-activity mutation suppresses the exocytosis and enhances the endocytosis of KCNQ3 at the AIS. Furthermore, our data on KCNQ3-ETD/AAA demonstrate that ankyrin-G is an important regulator for the exo/endocytosis process of KCNQ3.

**Figure 6.**
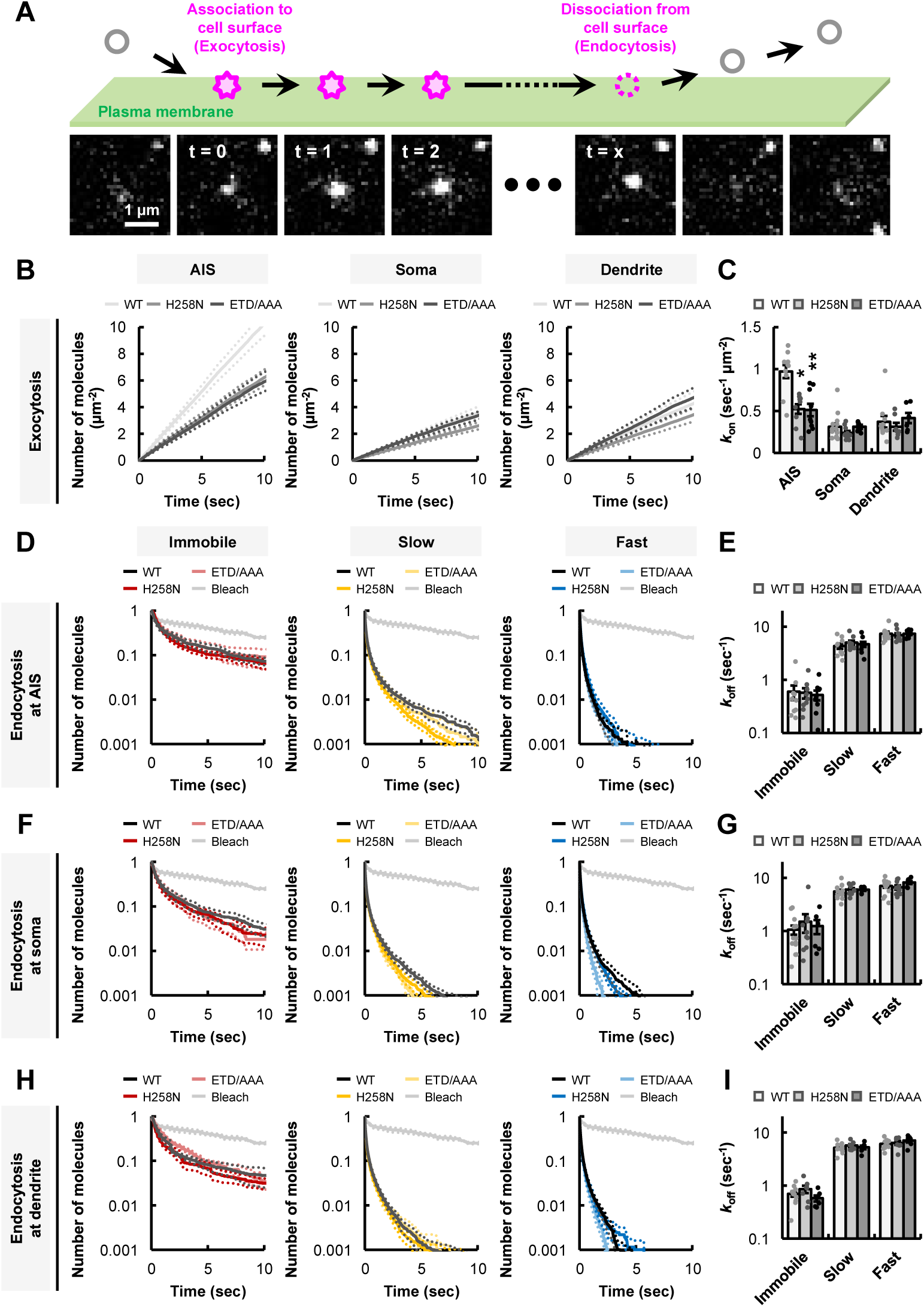
Exo/endocytosis of KCNQ2/3 at the AIS is altered by the low-activity mutation of KCNQ3. (A) A schematic diagram showing the exo/endocytosis events of Halo-TAC-KCNQ3 on the cell surface. Scale bar, 1 μm. (B and C) The association curves (B) and the *k*_on_ (C) of Halo-TAC-KCNQ3 on the cell surface at the AIS (WT, *n* = 10; H258N, *n* = 9; ETD/AAA, *n* = 9), the soma (WT, *n* = 14; H258N, *n* = 10; ETD/AAA, *n* = 6), and the dendrite (WT, *n* = 10; H258N, *n* = 7; ETD/AAA, *n* = 6). \**P* < 0.05 and \*\**P* < 0.01 (Steel-Dwass test, only comparisons with the “WT” group for the same region are depicted). (D-I) The dissociation curves (D, F, and H) and the *k*_off_ (E, G, and I) for three groups of Halo-TAC-KCNQ3 on the cell surface at the AIS (WT, *n* = 10; H258N, *n* = 9; ETD/AAA, *n* = 9), the soma (WT, *n* = 14; H258N, *n* = 10; ETD/AAA, *n* = 6), and the dendrite (WT, *n* = 10; H258N, *n* = 7; ETD/AAA, *n* = 6). The gray line shows the time course of bleaching and blinking at a rate of 0.150 ± 0.005 sec^-1^ (*n* = 3). *P* > 0.05 for all combinations among wild-type KCNQ3, KCNQ3-H258N and KCNQ3-ETD/AAA by Steel-Dwass test. Data are mean ± SE.

### Evidence that interaction between ankyrin-G and KCNQ2/3 is suppressed by the low-activity mutations of KCNQ3

The above results indicate that all of the three-dimensional dynamics of KCNQ3-H258N are closely similar to those of KCNQ3-ETD/AAA lacking ankyrin-G binding ability (Figures 5 and 6). This suggests that low-activity mutations may also affect the interaction between KCNQ2/3 and ankyrin-G. To verify this possibility, we analyzed the effect of ankyrin-G on the diffusion dynamics of wild-type and mutant KCNQ3 in HEK293T cells. The immobile state occupancy of wild-type KCNQ3 was increased without changing the diffusion coefficient upon co-expression of ankyrin-G (Figure 7A; Video S4). It was also found that ankyrin-G has a stronger effect on the diffusion of KCNQ3 than KCNQ2, although the diffusion coefficient of KCNQ2 was slightly but significantly decreased upon ankyrin-G co-expression (Figures S5A and S5B; Video S5). This is consistent with our results (Figure S1) and a previous report (Rasmussen et al. 2007) showing that KCNQ3, but not KCNQ2, functions as a dominant trafficking regulator of KCNQ2/3. In contrast to ankyrin-G, ankyrin-B did not affect the diffusion of KCNQ3, suggesting that KCNQ3 exhibits ankyrin isoform specificity (Figures S5C and S5D; Video S4). Co-expression of ankyrin-G did not increase the occupancy of the immobile state of KCNQ3-ETD/AAA, suggesting that KCNQ3 was immobilized on the cell surface by a direct interaction with ankyrin-G via the ABM of KCNQ3, even in HEK293T cells (Figure 7B; Video S6). However, the immobile state occupancies of the low-activity KCNQ3 mutants (KCNQ3-FDQ/RNE, KCNQ3-Y267C, and KCNQ3-H258N) were not affected upon ankyrin-G co-expression (Figures 7C-7E; Video S6). For simplicity, the ratio of the immobile state occupancy with and without ankyrin-G was calculated as an indicator of the ankyrin sensitivity of KCNQ3. The ankyrin sensitivities of the KCNQ3 mutants were significantly reduced compared to wild-type KCNQ3 (Figure 7F). Thus, our findings suggest that low-activity mutations of KCNQ3 interfere with the interaction between KCNQ2/3 and ankyrin-G at the AIS, causing a decrease in AIS selectivity.

**Figure 7.**
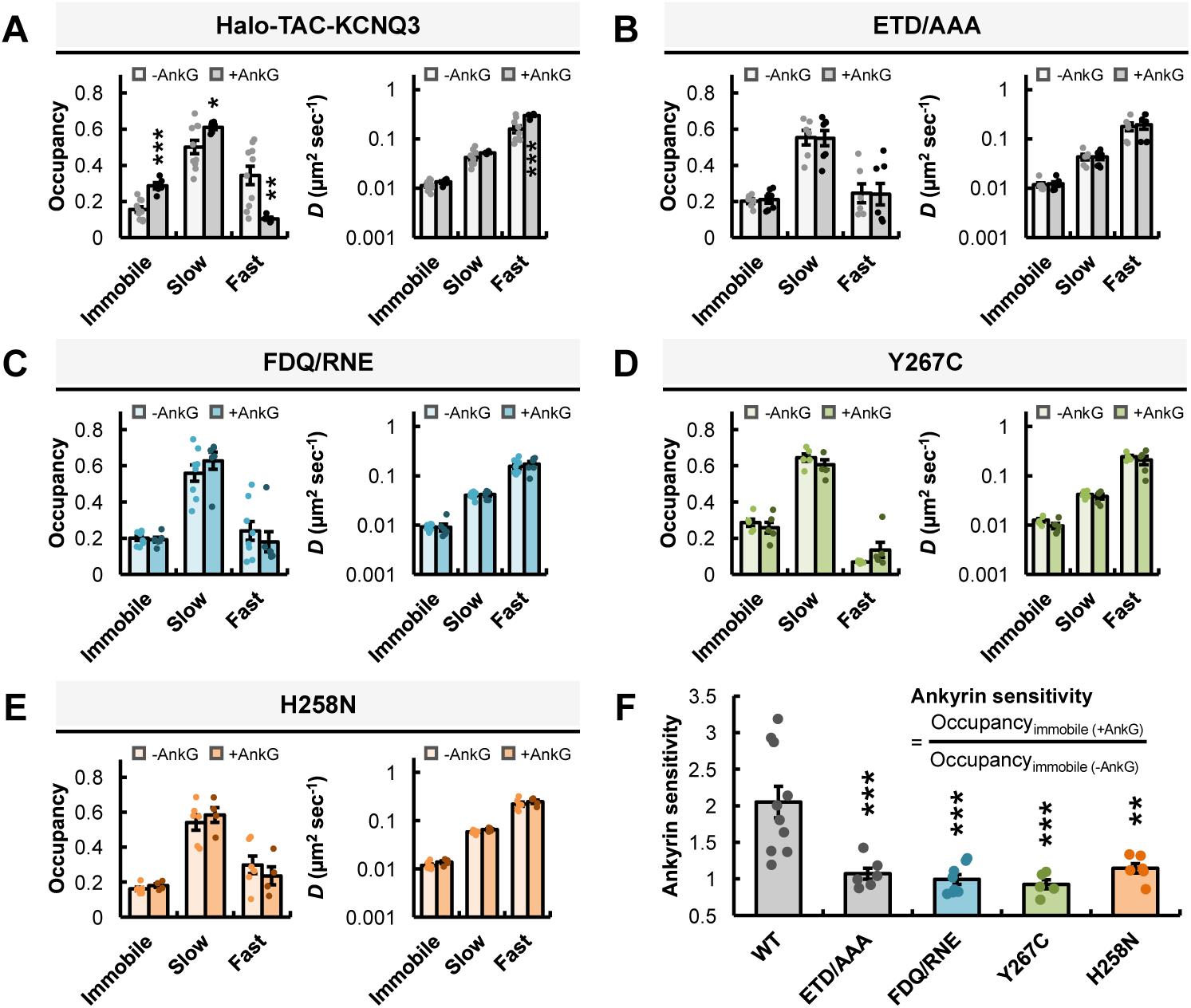
Evidence that interaction between ankyrin-G and KCNQ2/3 is suppressed by the low-activity mutations of KCNQ3. (A-E) Occupancy (left) and diffusion coefficient (right) for immobile, slow-diffusion, and fast-diffusion state of Halo-TAC-KCNQ3 in HEK293T cells with (A, WT, *n* = 5; B, ETD/AAA, *n* = 7; C, FDQ/RNE, *n* = 6; D, Y267C, *n* = 5; E, H258N, *n* = 4) or without (A, WT, *n* = 10; B, ETD/AAA, *n* = 6; C, FDQ/RNE, *n* = 8; D, Y267C, *n* = 5; E, H258N, *n* = 6) 270 kDa ankyrin-G. \**P* < 0.05, \*\**P* < 0.01, and \*\*\**P* < 0.001 (Welch’s *t*-test, comparisons were made with the “-AnkG” group for the same state). (F) Ankyrin sensitivity of Halo-TAC-KCNQ3 calculated as the ratio between the immobile state occupancy of with and without 270 kDa ankyrin-G (WT, *n* = 10; ETD/AAA, *n* = 6; FDQ/RNE, *n* = 8; Y267C, *n* = 5; H258N, *n* = 6). \*\**P* < 0.01 and \*\*\**P* < 0.001 (Dunnett’s test, comparisons were made with the “WT” column). Data are mean ± SE.

## Discussion

In this study, we employed multiple- and single-molecule imaging of Halo-TAC-KCNQ3 to directly visualize and comprehensively analyze the intricate trafficking process of KCNQ2/3 in neurons. We quantified the single-molecule dynamics of KCNQ2/3 at the AIS and somatodendritic region. It should be noted here that the quantified *k*_off_ and *k*_on_ were higher than expected, given the previously reported long-lifetime of AIS proteins on a day scale (Hedstrom, Ogawa, and Rasband 2008). Regarding the immobile molecules, their *k*_off_ and *k*_on_ were affected by bleaching or blinking of the fluorescent ligand (TMR) at a rate of 0.150 ± 0.005 sec^-1^ (Figure 6), and were overestimated compared to the true values. Regarding the slow- and fast-diffusing molecules, their *k*_off_ and *k*_on_ were not affected by bleaching or blinking (Figure 6). Instead, it is possible that they reflect the recycling process where vesicles containing Halo-TAC-KCNQ3 transiently fuse with or simply contact the plasma membrane and then immediately return to the cytoplasm, supporting the necessity of molecular classification based on their dynamics. Although the exact lifetime of immobile KCNQ2/3 could not be measured, we found that their diffusion is confined to the radii of 33 nm (AIS), 172 nm (soma), and 142 nm (dendrite), regardless of their lifetime (Figure S3). The diffusion distances of slow- and fast-diffusing molecules during their lifetime can be calculated as the square root of 4*D*/*k*_off_ (Figures 5 and 6): for slow-diffusing molecules, 153 nm (AIS), 144 nm (soma), and 140 nm (dendrite); for fast-diffusing molecules, 242 nm (AIS), 226 nm (soma), and 238 nm (dendrite). Since the estimated diffusion distances of KCNQ2/3 are orders of magnitude smaller than the spatial scale of neurons, the effect of lateral diffusion on the AIS localization of KCNQ2/3 in mature neurons is assumed to be small. In contrast, the surface density of KCNQ2/3 determined by exo/endocytosis can be calculated as *k*_on_/*k*_off_ (Figure 6), and the AIS to dendrite molecular density ratio based on the *k*_on_ and *k*_off_ was 3.433. This indicates that exo/endocytosis alone is enough to account for the AIS localization of KCNQ2/3 without considering lateral diffusion, demonstrating the detailed mechanism for spatial pattern formation of KCNQ2/3 in mature neurons. Taken together with a previous report on immature neurons (Ghosh et al. 2020), our results suggest that the mechanism regulating AIS targeting of KCNQ2/3 switches from a lateral diffusion-dependent manner to an exo/endocytosis-dependent manner during the neuronal development from immaturity to maturity.

All the trafficking pathways of KCNQ3 are regulated by ankyrin-G. Single-molecule imaging analysis revealed that the ETD/AAA mutation in the ABM of KCNQ3 suppressed the exocytosis, increased the lateral diffusivity, and enhanced the endocytosis of KCNQ2/3 at the AIS, but not at the somatodendritic region (Figures 5 and 6; Table S1). In particular, the AIS to dendrite molecular density ratio based on the *k*_on_ and *k*_off_ of KCNQ3-ETD/AAA was 1.442, suggesting that exo/endocytosis sufficiently accounts for the less AIS selectivity of KCNQ3-ETD/AAA lacking ankyrin-G binding ability. Ankyrin-G is well known as the major scaffold protein at the AIS which primarily defines the AIS assembly and interacts with various AIS proteins, including KCNQ2/3 (Jenkins and Bennett 2001). Ankyrin-G not only restricts the lateral diffusion (Benned-Jensen et al. 2016), but also affects the exo/endocytosis of AIS proteins. Regarding the exocytosis, the interaction with ankyrin-G is necessary for the recruitment of Nav1.6 to the AIS region via direct exocytosis in neurons (Akin et al. 2015). Regarding the endocytosis, the inhibitory effects on the endocytosis of AIS proteins by interaction with ankyrin-G have been reported (Fréal et al. 2019; Torii et al. 2020; Eichel et al. 2022). Given the multifaceted role of ankyrin-G in KCNQ2/3 trafficking regulation, the interaction with ankyrin-G can consistently explain the clear correlation between the diffusivity and lifetime of KCNQ2/3, and the differences in all parameters of spatial dynamics between wild-type KCNQ3 and KCNQ3-ETD/AAA at the AIS (Figures 5 and 6; Table S1). It should be noted here that the physical properties of the slow-diffusion state of KCNQ3 were not changed by the ETD/AAA mutation. Intriguingly, the confinement radius of the slow-diffusion state of KCNQ3 at the AIS (70 nm) was similar to the half-width between actin rings at the AIS (95 nm) (Leterrier et al. 2015), but was extended to 257 nm at the dendrite where similar actin rings form (D’Este et al. 2015) (Figure S3). These results suggest that the slow-diffusion state of KCNQ3 is not defined by an ankyrin G-dependent mechanism, but rather by an ankyrin-G independent one such as the “fence and picket” model of diffusion barriers at the AIS (Winckler, Forscher, and Mellman 1999; Nakada et al. 2003). Notably, the binding affinity of KCNQ3 for ankyrin-B is lower than that for ankyrin-G (Figure S5). This demonstrates why KCNQ3 asymmetrically localizes at the AIS, even though ankyrin-B accumulates in the distal axon and dendrite in a manner that is mutually exclusive to ankyrin-G (Galiano et al. 2012; Lorenzo et al. 2014).

Most surprisingly, mutations in the activity regulatory sites of KCNQ3 inhibited AIS targeting of KCNQ2/3 in a channel functionality-dependent manner (Figure 2). The single-molecule dynamics of the KCNQ3-H258N were closely similar to that of KCNQ3-ETD/AAA (Figures 5 and 6; Table S1), and the AIS to dendrite molecular density ratio based on the *k*_on_ and *k*_off_ of KCNQ3-H258N was 1.882. Consistent with the observations in neurons, ankyrin-G did not restrict the lateral diffusion of low-activity KCNQ3 mutants (KCNQ3-FDQ/RNE, KCNQ3-Y267C, and KCNQ3-H258N) in HEK293T cells (Figure 7). These results suggest that low-activity KCNQ3 mutants have a lower binding affinity to ankyrin-G compared to wild-type KCNQ3, resulting in the observed differences in the spatial dynamics between wild-type and low-activity mutant KCNQ3. In addition to the ankyrin-G dependent mechanism, theoretically, several other mechanisms could also potentially account for the less AIS selectivity observed in low-activity KCNQ3 mutants. These include: 1) a reduction in the cytoplasmic pool of KCNQ2/3; 2) a disruption of heteromeric KCNQ2/3 complex formation; and 3) a reorganization of the AIS triggered by increased neuronal excitability. However, our data show that the protein expression of KCNQ3 is not affected by mutations in the activity regulatory sites (Figure 2), that low-activity KCNQ3 mutants have the ability to form KCNQ2/3 complexes (Figure 3), and that the expressions of the low-activity KCNQ3 mutants does not alter the neuronal excitability and AIS structure (Figure 4), ruling out the above hypothesis. In summary, we revealed that mutations in the activity regulatory sites of KCNQ3 impact not only the functionality but also the trafficking process of KCNQ2/3 via interaction with ankyrin-G, thereby reducing the AIS selectivity.

Pathologically, mutations of KCNQ2/3 have been linked to various neurological diseases, including epilepsy (Biervert et al. 1998; Orhan et al. 2014; Nappi et al. 2020). Intriguingly, clusters of epilepsy-related mutations that cause trafficking abnormalities reported to date are predominantly distributed in the PD (Chung, Jan, and Jan 2006; Abidi et al. 2015) and cytoplasmic C-terminal region (Chung, Jan, and Jan 2006; Alaimo et al. 2009; Kim et al. 2018) of KCNQ2/3. Here, we identified trafficking-related sites in regions other than the PD and C-terminal regions (Figure 2). Notably, some of the activity regulatory sites of mouse KCNQ3 that we examined (R243, H258, Y267, and R365) correspond to mutated residues found in disease-related variants of human KCNQ3, namely R242Q/W, H257Y, Y266C (Symonds et al. 2019), and R364H/C (ClinVar accession ID: VCV000934080.14; VCV001376796.5; VCV000937869.5; VCV000205964.2; VCV000934997.12; VCV000656938.6). This indicates that the mutations of these residues impair both functionality and trafficking of KCNQ2/3, thus demonstrating the pathogenic effects of these mutations on epileptic diseases.

The coupling between functionality and trafficking can hold pathophysiological significance in patients who carry disease mutation of KCNQ3 heterozygously. If low-activity KCNQ2/3 containing mutant KCNQ3 were normally transported to the cell surface at the AIS, the mutant KCNQ2/3 could occupy ankyrin-G and competitively inhibit surface expression of normal KCNQ2/3. This would cause attenuation of KCNQ2/3-current and thus increased neural excitability. Our data show that the AIS surface expression of low-activity KCNQ2/3 is inhibited (Figures 2 and S1). The low trafficking efficiency of mutant KCNQ2/3 can contribute to maintain a high occupancy of normal KCNQ2/3 on the AIS surface and suppresses the KCNQ2/3-current attenuation, thereby serving as a safety mechanism. This inference is well consistent with our observation that expression of the KCNQ3-H258N mutant did not significantly alter intrinsic neuronal excitability (Figure 4). In this context, the functionality-trafficking coupling of KCNQ2/3 could be considered as a protein quality control mechanism on the neuronal cell surface. However, in human patients who carry disease mutation homozygously, the functionality-trafficking coupling can lead to severe symptoms by significantly reducing the number of KCNQ2/3 molecules on the cell surface. This hypothesis could explain why mutations in KCNQ3 result in more severe disease states and why disease-related mutations in KCNQ3 are rarer compared to KCNQ2 (Nappi et al. 2020). Furthermore, this coupling presents a significant challenge in drug screening for the epileptic channelopathies. This is because conventional channel openers such as retigabine cannot exert their functions, if the target molecules are not properly trafficked and do not exist on the cell surface. This could lead to drug-resistant epilepsy, underscoring the importance of considering the spatial dynamics of target molecules in the drug screening. Taken together, our findings not only deepen the understanding of KCNQ2/3 regulation mechanism but also provide a profound insight into pathogenesis and therapeutic strategies for neurological disorders caused by the defects in the KCNQ2/3.

## Acknowledgements

We thank Vann Bennett (Duke University, US) for the constructs of EGFP-fused 270 kDa ankyrin-G and EGFP-fused 220 kDa ankyrin-B. We also thank Yuko Fukata (National Institute for Physiological Sciences, Japan) and Koichiro Irie (Osaka University, Japan) for their invaluable technical advice regarding the materials and methods used in the primary culture of hippocampal neurons. This research was supported by Japan Society for the Promotion of Science (JSPS) KAKENHI Grant (JP20K22631 and JP22K15373 to D.Y.; 21K19350 and 23K24066 to Y.O.); The Protein Research Foundation (to D.Y.); The Uehara Memorial Foundation (to D.Y.); The Mitsubishi Foundation (to Y.O.); and ONO Medical Research Foundation (to Y.O.).

## Author Contributions

D.Y. designed and conducted all the experiments, and wrote the paper. Y.O. designed the experiments and wrote the paper.

## Methods

### DNA constructs

Mouse KCNQ2 (852 amino acids; UniProt accession number: B7ZBW1) and mouse KCNQ3 (873 amino acids; UniProt accession number: Q8K3F6) were cloned into the pCS5+ vector for expression in mammalian cells. HaloTag (Promega) and a single transmembrane region from human IL2RA (UniProt accession number: P01589) were fused to the N-terminus of KCNQ2 or KCNQ3 using the In-Fusion HD Cloning Kit (Takara). Site-directed mutations were introduced into KCNQ2 and KCNQ3 by standard PCR techniques using PrimeSTAR Max DNA Polymerase (Takara). For whole-cell patch clamp recording of KCNQ3 channels in HEK293T cells, the A316T mutation was introduced to facilitate efficient trafficking and functional expression of the homomeric KCNQ3 channel complex (Etxeberria et al. 2004). The pl-Synapsin-YFP-NavII-III construct (AIS marker) was obtained from Addgene (Plasmid #91246), and the fluorescent tag was replaced with EGFP. EGFP-fused 270 kDa ankyrin-G and EGFP-fused 220 kDa ankyrin-B were gifts from Vann Bennett (Duke University, Durham, US). The DNA sequences of all constructs were confirmed through Sanger sequencing.

### Culture of HEK293T cells

HEK293T cells were cultured in DMEM (10% FBS, 0.1% penicillin/streptomycin) at 37°C, 5% CO_2_. The cells were passaged every 2-3 days. For the whole-cell patch clamp experiment, a total of 300 ng of plasmid was transfected into the cells on a 35-mm dish using Lipofectamine 2000 (Invitrogen). For the measurement of the homomeric KCNQ2 or KCNQ3T, 200 ng of KCNQ2 or Halo-TAC-KCNQ3T (wild-type or the mutants) was co-transfected with 100 ng of EGFP. For the measurement of the heteromeric KCNQ2/3 complex, 100 ng each of KCNQ2, Halo-TAC-KCNQ3 (wild-type or the mutants), and EGFP were co-transfected. In both cases, whole-cell patch clamp recordings were performed 24 hours post-transfection. For single-molecule imaging under TIRFM, 200 ng of Halo-TAC-KCNQ3 (wild-type or the mutants) or Halo-TAC-KCNQ2 was transfected. When ankyrin-G or ankyrin-B was co-expressed in Figures 7 and S5, 800 ng of EGFP fused 270 kDa ankyrin-G or EGFP fused 220 kDa ankyrin-B was co-transfected and imaging was performed 48 hours post-transfection. The imaging data in Figures S2A-S2C were obtained 24 hours after co-transfection with 100 ng of EGFP.

### Primary culture of hippocampal neurons

The hippocampi were dissected from the brain of fetal C57BL/6 mice (E17). The sex of the mice was not considered for the primary culture of hippocampal neurons, and the tissues derived from both male and female mice were mixed. For digestion, the tissues were washed with PBS on ice, and then treated with a 3 ml PBS solution additionally containing 25 mg glucose, 16.5 mg papain (164-00172, FUJIFILM), 1 mg L-cysteine (10309-41, Nacalai Tesque), 1 mg BSA (A9418-50G, SIGMA), and 1,500 U DNase I Type II (D4527-40KU, SIGMA) for 20 min at 37°C. Subsequently, the digested tissues were washed with 5 mL of DMEM (10% FBS) to halt the enzymatic reaction of papain. The supernatant was subsequently removed, and the tissues were mechanically triturated by pipetting in 1 ml of HBSS (17460-15, Nacalai Tesque) containing 3 mg MgSO_4_·7H_2_O and 500 U DNase I Type II. The suspension of dissociated neurons was collected into a new tube containing 5 mL of DMEM (10% FBS) and centrifuged at 1,000 × g for 5 min at 4°C. After removing the supernatant, the pellet was resuspended in Neurobasal medium (Gibco) supplemented with GlutaMAX (Sigma) and B-27 (Gibco). The cell density was adjusted to 2 x 10^5^ cells/ml, and the 1 ml suspension of neurons was plated onto glass coverslips (Matsunami) coated with Poly-L-Lysine (28360-14, Nacalai Tesque) and collagen (354236, CORNING) in 24-well plates. After plating, half of the medium was replaced with fresh medium twice a week. At 3 DIV, 2 μM Ara-C (030-11951, FUJIFILM) was added to the medium to prevent the proliferation of non-neuronal cells. At 7 DIV, a total of 1.2 μg of plasmid DNA was transfected into the neurons using Lipofectamine 2000. Each plasmid DNA (e.g. Halo-TAC-KCNQ3, KCNQ2, EGFP-Nav1.2 II-III linker) was transfected at the same amount (400 ng).

### Electrophysiology

Whole-cell configuration patch clamp recordings were conducted in HEK293T cells and cultured neurons using an amplifier (EPC 800, HEKA) and a digitizer (Digidata 1550A, Molecular Devices). The signals were sampled at 20 kHz and filtered at 5 kHz. The junction potential was not adjusted. The series resistance compensation was not conducted. Microelectrodes were made from borosilicate glass capillaries (B150-86-10, Sutter Instrument) using a puller (P-97, Sutter Instrument).

For the recording of HEK293T cells, the extracellular solution was composed of 160 mM NaCl, 5 mM KCl, 2 mM CaCl_2_, 1 mM MgCl_2_, and 10 mM HEPES (pH 7.4 adjusted with NaOH). The microelectrodes with a resistance of 2-5 MΩ were filled with the intracellular solution composed of 160 mM KCl, 5 mM MgCl_2_, 2 mM Na_2_ATP and 10 mM HEPES (pH 7.4 adjusted with KOH). The current was examined by holding the membrane potential at -80 mV and applying a 300 msec depolarizing pulse from -80 mV to 60 mV in 10 mV increments every 3 sec. The conductance-voltage curve (*G*-*V*) was obtained by converting the current value at each voltage step to conductance using the equation *G* = *I*/(*V*-*V*_rev_), where *G* is the conductance, *I* is the peak current, *V* is the command pulse potential, and *V*_rev_ is the reversal potential. *V*_rev_ was calculated using the Goldman-Hodgkin-Katz voltage equation, based on the relative Na^+^ permeability compared to that of K^+^ in KCNQ2 and KCNQ3T as reported previously (Manville and Abbott 2019). The *G*-*V* curve was fitted with the Boltzmann equation *G*/*G*_max_ = 1/(1+exp((*V*_1/2_-*V*)/*k*)), where *G*_max_ is the maximum conductance, *V*_1/2_ is the voltage required to activate half-maximum conductance, and *k* is the slope coefficient of the *G*-*V* curve. The activation time constant (*τ*) was determined by fitting a single exponential function to the KCNQ3T- or KCNQ2/3-current, 10 msec after the onset of the depolarizing voltage pulse to avoid the sigmoidal activation phase. The voltage dependence of *τ* was assessed by fitting the *τ*-*V* plot with the equation *τ* = *A*exp(-*V*/*x*)+*τ*_0_, where *τ* is the activation time constant, *V* is the membrane potential, *A* is the amplitude factor, *τ*_0_ is the minimum value of *τ*, *x* is the slope coefficient. For the recording of primary cultured neurons (10-13 DIV), the extracellular solution was composed of 160 mM NaCl, 2.5 mM KCl, 10 mM HEPES, 10 mM glucose, 1.2 mM MgCl_2_, and 1.8 mM CaCl_2_ (pH 7.3 adjusted with NaOH). The microelectrodes with a resistance of 2-6 MΩ were filled with the intracellular solution composed of 135 mM KCl, 2 mM EGTA, 1.1 mM CaCl_2_, 10 mM HEPES, and 5 mM glucose (pH 7.25 adjusted with KOH). The data were processed and analyzed using Clampfit 10.7 (Molecular Devices).

### Subcellular localization imaging under confocal microscopy

The HaloTag-fused proteins expressed in neurons (10 DIV) placed on coverslips were fluorescently labeled with 1 µM AF660 (Promega) in medium for 15 min at 37°C, 5% CO_2_. The HaloTag-fused proteins were subsequently labeled with 1 µM TMR (Promega) in medium for 15 min at 37°C, 5% CO_2_. The neurons were then washed three times with 1 ml of an artificial cerebrospinal fluid solution (aCSF; 125 mM NaCl, 5 mM KCl, 1 mM MgCl_2_, 2 mM CaCl_2_, 10 mM glucose, and 10 mM HEPES, pH 7.4 adjusted with NaOH). The coverslip was mounted in a magnetic chamber (CM-B18-1, Live Cell Instrument).

The spatial distribution of fluorescent molecules in neurons was imaged using a confocal laser scanning microscope (FV3000, Olympus) equipped with an oil immersion objective lens (UPLSAPO60X, 60×, Oil, NA 1.42, Olympus). The pixel size was 414 nm. All images were processed and analyzed with Fiji (Schindelin et al. 2012). Images were acquired by merging z-stacks with a step size of 0.41 μm. Prior to the analysis, a Gaussian filter (2 pixels) was applied. The profile of fluorescence intensity along the axon was plotted from the edge of the soma. The mean value of the AF660 signal was obtained at axons 20-40 μm and at dendrites 0-20 μm from the edge of the soma as the fluorescence intensity at the AIS and dendrite, respectively (for data shown in Figures 1-3; Figure S1). The mean value of the TMR signal was obtained at axons 0-20 μm from the edge of the soma (for data shown in Figures 1-3; Figure S1). To analyze the morphological properties (length and start position) of the spatial distribution of molecules, the profile of fluorescence intensity along axons was min-max normalized. The threshold value was set at 0.5 (half maximum), and the length and start position of the longest contiguous region above the threshold were quantified.

### Single-molecule imaging under TIRFM

For the measurement of HEK293T cells, cells transfected with the specified plasmid DNA were plated on coverslips coated with collagen (TMTCC-050, TOYOBO) a few hours before measurement. The subsequent fluorescent labeling and imaging were conducted after the cells had sufficiently adhered to the coverslips. The cells were labeled with 2.5 pM TMR for 15 min at 37°C, 5% CO_2_. The cells were then washed three times with 1 ml of HBSS and incubated for a further 15 min at 37°C, 5% CO_2_. For the measurement of cultured neurons (10 DIV), the neurons were labeled with 2.5 pM TMR for 15 min at 37°C, 5% CO_2_. The neurons were then washed three times with 1 ml of an aCSF and incubated for a further 15 min at 37°C, 5% CO_2_. The coverslip was mounted in a metal chamber (for HEK293T cells; Attofluor Cell Chamber, Invitrogen) or a magnetic chamber (for neurons; CM-B18-1, Live Cell Instrument) and then additionally washed two times with 0.5 ml of HBSS (for HEK293T cells) or aCSF (for neurons) before imaging under TIRFM.

Single-molecule imaging was performed using an inverted microscope (IX71, Olympus) equipped with an oil immersion objective lens (UPLAPO100XOHR, 100×, Oil, N.A. 1.50, Olympus) and a 1.6× intermediate magnification lens, resulting in a total magnification of 160× (the pixel size was 100 nm). TMR-labeled molecules near the bottom cell surface were excited with 532 nm lasers (OBIS 532 nm, 50 mW, COHERENT) through the objective lens by a dichroic mirror (ZT532/640rpc, CHROMA). The images were captured by an electron-multiplying charge-coupled device (EM-CCD) camera (ImagEM, Hamamatsu) after passing through a single band pass filter (FF01-593/40-25, Semrock). The penetration depth (*d*) of the TIRFM evanescent field was estimated as follows,

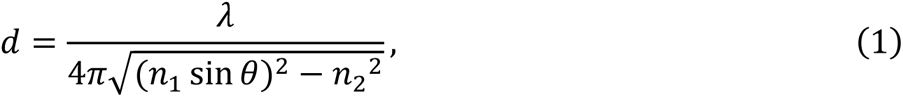

where *λ* is the wavelength of the laser light (532 nm), *n*_1_ is the refractive index of oil and glass (1.52), *n*_2_ is the refractive index of water (1.33), and *θ* is the angle of incidence of the laser light (0.9 rad). Given these conditions, the penetration depth (*d*) of the evanescent field was estimated to be 101.4 nm. Fluorescent images (256 × 512 pixels) were acquired using image software (MetaMorph, Molecular Devices) with an exposure time of 33 msec and an electron-multiplying (EM) gain setting of 1200. All measurements were performed at room temperature.

### Single-particle tracking

Acquired 16-bit TIFF files were processed using Fiji. As a pre-processing measure, background subtraction was performed using a rolling ball algorithm with a radius of 5 pixels, followed by Gaussian filtering (0.5 pixels). The region of interest (ROI) was then manually defined. Using the Fiji plug-in TrackMate (Tinevez et al. 2017), the positions of all fluorescent spots were automatically determined across all frames of the movie. A Laplacian of Gaussian (LoG) detector was utilized for the detection of these bright spots. Fluorescent signals exhibiting abnormal characteristics were excluded by setting thresholds for both the size and brightness of the spots (Estimated blob diameter = 4.5 pixels; Threshold = 500). Subsequently, a simple Linear Assignment Problem (LAP) tracker was employed to link particles between frames. Fluorescent spots in adjacent frames were considered to represent the same fluorescent molecule if the distance between them fell within the typical value for molecular diffusion (Linking max distance = 2.5 pixels; Gap-closing max distance = 2.5 pixels; Gap-closing max frame gap = 1). All molecules with the lifetime exceeding 2 frames were included in the subsequent data analysis.

### Analysis of single-molecule dynamics

Initially, the number of diffusion states of KCNQ3 at the AIS was evaluated as previously described (Yoshioka et al. 2020). Specifically, the probability density distribution of displacements (*r*) that a molecule moved during exposure time (Δ*t* = 33 msec) was fitted with the sum of two-dimensional diffusion equations:

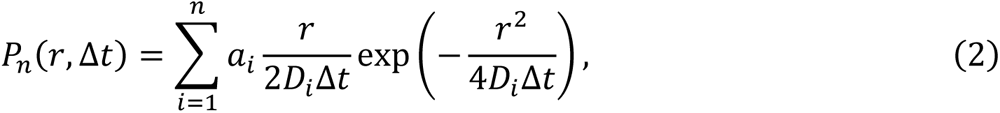

where *n* represents the total number of diffusion states in the assumed model, which was manually set. *D*_i_ and *a*_i_ denote the diffusion coefficient and the occupancy of the *i*-th diffusion state in the *n*-state model, respectively. Optimal parameters (*D*_i_ and *a*_i_) were estimated using the maximum likelihood estimation method based on the Expectation-Maximization (E-M) algorithm. In order to determine the optimal number of states (*n*) for explaining the experimental data, the Akaike Information Criterion (AIC) was utilized:

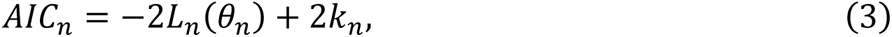

where *θ*_n_ denotes the matrix of parameters (*D*_i_, *a*_i_) to be estimated in the *n*-state model, and *L*_n_ denotes the log likelihood expressed as:

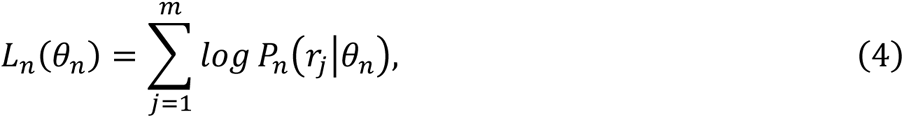

where *m* represents the total number of all *r* obtained from the single-molecule trajectory data, and *r*_j_ denotes the *j*-th *r* (*j* = 1, 2, … *m*). Based on this analysis, the 3-state model was determined to be the most suitable for elucidating the actual diffusion dynamics of wild-type KCNQ3 at the AIS (Figures S2G-S2I). Therefore, the 3-state model was uniformly used for the analysis of all single-molecule imaging data in this study.

Next, single-molecule trajectories were classified into 3 states using a Hidden Markov Model (HMM). The Baum-Welch algorithm (Baum 1972) was employed to estimate the initial probability *π*_i_ (0 ≤ *π*_i_ ≤ 1 and ∑_i_ *π*_i_ = 1), diffusion coefficient *D*_i_, and state transition probability matrix *A* (0 ≤ *A*_ij_ ≤ 1 and ∑_j_ *A*_ij_ = 1; *A*_ij_ denotes the transition probability from the *i*-th state to the *j*-th state) for each state (*i*, *j* = 1, 2, 3) (Figures 5 and S4A-S4C). The probability of displacement during exposure time (Δ*t* = 33 msec) was described using the two-dimensional diffusion equation,

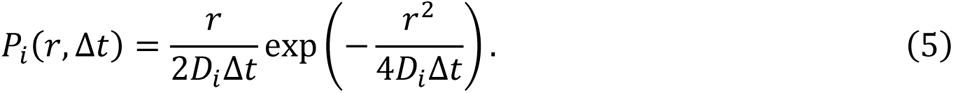

The occupancy of each diffusion state in the total trajectories was calculated as the mean probabilities of each state at each time point. Additionally, the Viterbi algorithm (Viterbi 1967) was utilized to classify the states of the molecules at each time point.

The ensemble Mean Squared Displacement (MSD) was calculated using the trajectories of molecules in each diffusion state. The ensemble MSD curve was fitted with the circular confinement diffusion model presented as follows:

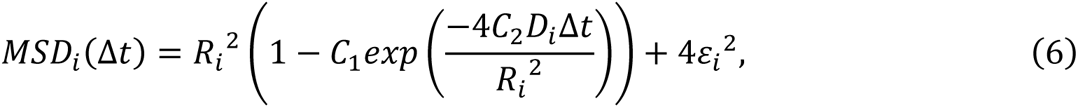

where Δ*t* represents the time lag, *D*_i_, *R*_i_, and *ε*_i_ denote the diffusion coefficient, confinement radius, and position error for each state (*i* = 1, 2, 3) respectively (Figure S3). Constants *C*_1_ and *C*_2_ were set to 0.99 and 0.85, assuming a circular confinement region (Michael J. Saxton and Jacobson 1997; Pinaud et al. 2009). This accounted for the effect of position error, resulting in diffusion coefficients closer to the true values.

To investigate the exocytosis process, an association plot was generated by accumulating the density of newly appearing molecules on the cell surface at each time point. This association plot depicts the exocytosis kinetics of molecules on the cell surface. The exocytosis rate constant (*k*_on_) was quantified as the slope of the association plot. It should be noted that the *k*_on_ includes the factor of molecular density in the cytoplasmic region, although no significant difference in molecular density in the cytoplasmic region (TMR signal) was detected between the wild-type and mutant KCNQ3 (KCNQ3-ETD/AAA and KCNQ3-H258N) (Figures 1O and 2H). To investigate the endocytosis process, molecules were initially categorized into immobile, slow-, and fast-diffusing groups. This classification was based on the diffusion state that maximized their occupancy in the trajectories (Figures S4D-S4L). As previously described (Yoshioka et al. 2020), the lifetime of each molecule on the cell surface was measured for each group. A dissociation plot was generated from the cumulative distribution of these lifetimes, illustrating the endocytosis kinetics of molecules from the cell surface. The endocytosis rate constant (*k*_off_) was quantified as the inverse of the average lifetime (Figure 6). The fraction of immobile, slow-, and fast-diffusing molecules was calculated using the number of each classified molecule appearing per 10 µm² over 15 sec in each neuron (Table S1).

### Statistics and reproducibility

Values are expressed as the mean ± SE containing a specified number of independent cells. Statistical significance of differences between groups was determined using paired *t*-test or Welch’s *t*-test. Statistical significance of differences in the fraction of classified molecules between groups was determined using Chi-square test and subsequent residual analysis. Steel-Dwass test was used for comparisons among three groups. Dunnett’s test was used for comparisons among more than three groups. Significance levels are \**P* < 0.05, \*\**P* < 0.01, and \*\*\**P* < 0.001. Details regarding the number of biological replicates and the assays are provided in the figure legends.

## Data availability

The source data for this paper and the code used for the analysis are available from the corresponding authors upon reasonable request.

