## Supplementary Information for "Coupling Between Functionality and Trafficking to the Axon Initial Segment in KCNQ2/3 K^+^ Channels"

### Supplementary Figures

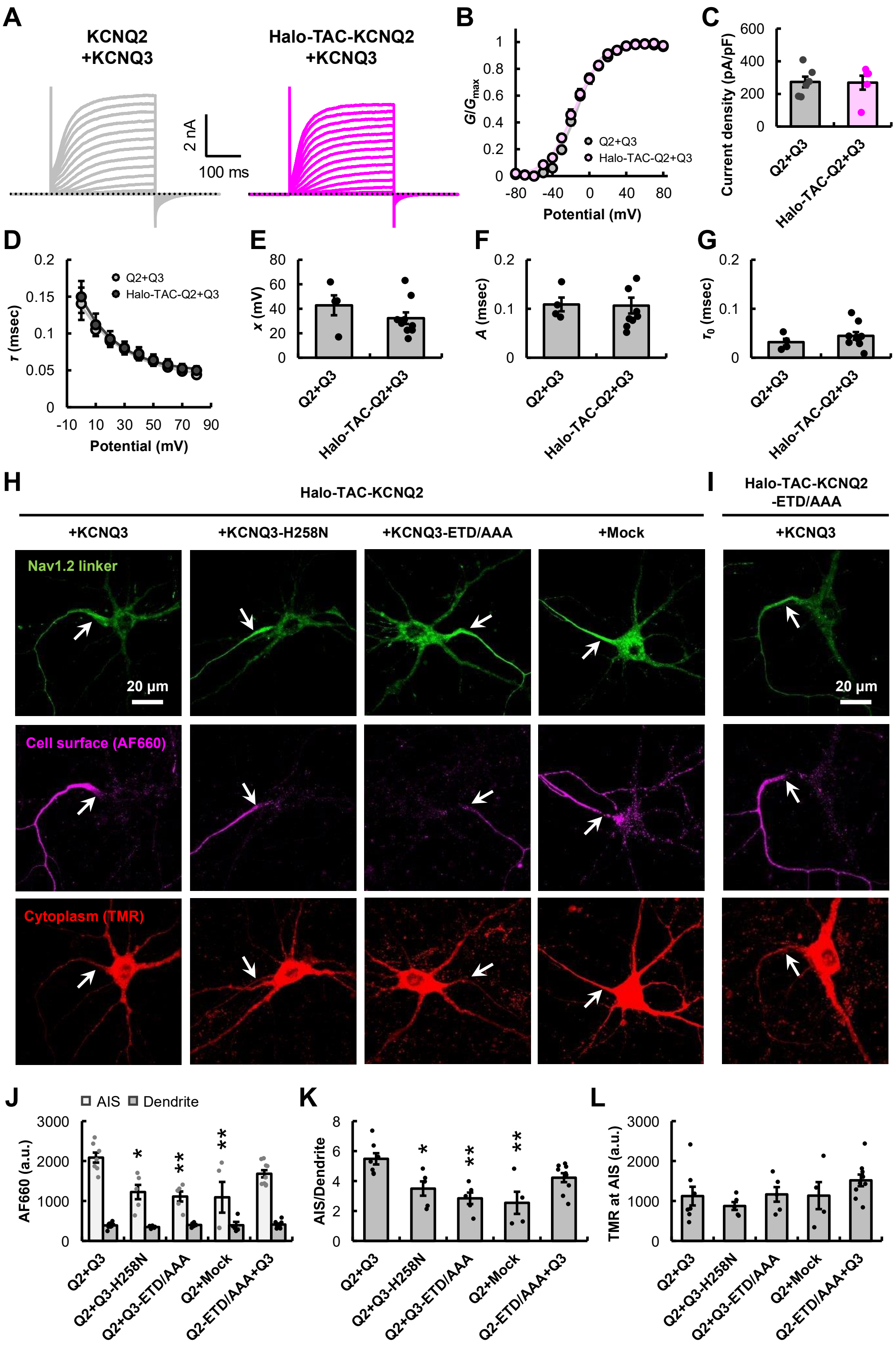


### Figure S1

**Figure S1: Dominant-negative effect of mutant KCNQ3 on the AIS localization of Halo-TAC-KCNQ2 in mouse hippocampal neurons.**

(A) Representative current traces of KCNQ2+KCNQ3 (left) or Halo-TAC-KCNQ2+KCNQ3 (right) obtained by whole-cell patch-clamp recordings in HEK293T cells. (B) *G*-*V* curves of KCNQ2+KCNQ3 (Q2+Q3) and Halo-TAC-KCNQ2+KCNQ3 (Halo-TAC-Q2+Q3). $V_{1/2}$ for KCNQ2+KCNQ3: -12.22 ± 1.58 mV ($n = 6$) and for Halo-TAC-KCNQ2+KCNQ3: -15.44 ± 1.61 mV ($n = 5$). $P = 0.230$ (Welch's *t*-test). (C) Current densities at 80 mV for KCNQ2+KCNQ3 (Q2+Q3, $n = 6$) and Halo-TAC-KCNQ2+KCNQ3 (Halo-TAC-Q2+Q3, $n = 5$). $P = 0.943$ (Welch's *t*-test). (D) Activation time constant ($\tau$) of KCNQ2+KCNQ3 ($n = 4$) and Halo-TAC-KCNQ2+KCNQ3 ($n = 9$) at different potentials. (E-G) The quantified parameters: $x$ (E, $P = 0.370$), $A$ (F, $P = 0.924$), and $\tau_0$ (G, $P = 0.290$). *P*-values were obtained by Welch's *t*-test. (H) Representative confocal microscopic images of primary cultured hippocampal neurons expressing EGFP-Nav1.2 II-III linker (green) and Halo-TAC-KCNQ2 (magenta and red) with wild-type KCNQ3, KCNQ3-H258N, KCNQ3-ETD/AAA, or empty vector. Halo-TAC-KCNQ2 was labeled with AF660 (magenta) and TMR (red). The arrowhead indicates the position of AIS. Scale bar, 20 µm. (I) Representative confocal microscopic images of primary cultured hippocampal neurons expressing EGFP-Nav1.2 II-III (green) and Halo-TAC-KCNQ2-ETD/AAA (magenta and red) with wild-type KCNQ3. Halo-TAC-KCNQ2-ETD/AAA was labeled with AF660 (magenta) and TMR (red). The arrowhead indicates the position of AIS. Scale bar, 20 µm. (J-L) Average AF660 intensity at the AIS and dendrite (J), ratio of AF660 intensity between AIS and dendrite (K), TMR intensity at the AIS (L) of neurons expressing Halo-TAC-KCNQ2 with wild-type KCNQ3 (Q2+Q3, $n = 7$), KCNQ3-H258N (Q2+Q3-H258N, $n = 5$), KCNQ3-ETD/AAA (Q2+Q3-ETD/AAA, $n = 5$), or empty vector (Q2+Mock, $n = 4$), and neurons expressing Halo-TAC-KCNQ2-ETD/AAA with wild-type KCNQ3 (Q2-ETD/AAA+Q3, $n = 9$). $*P < 0.05$ and $**P < 0.01$ (Dunnett's test; comparisons were made with the "Q2+Q3" column for the same group). Data are mean ± SE. These data indicate that the low-activity mutant KCNQ3 (KCNQ3-H258N) significantly inhibits the AIS targeting of KCNQ2 in a dominant-negative manner, and that KCNQ3, rather than KCNQ2, is the primary regulator of KCNQ2/3 trafficking.

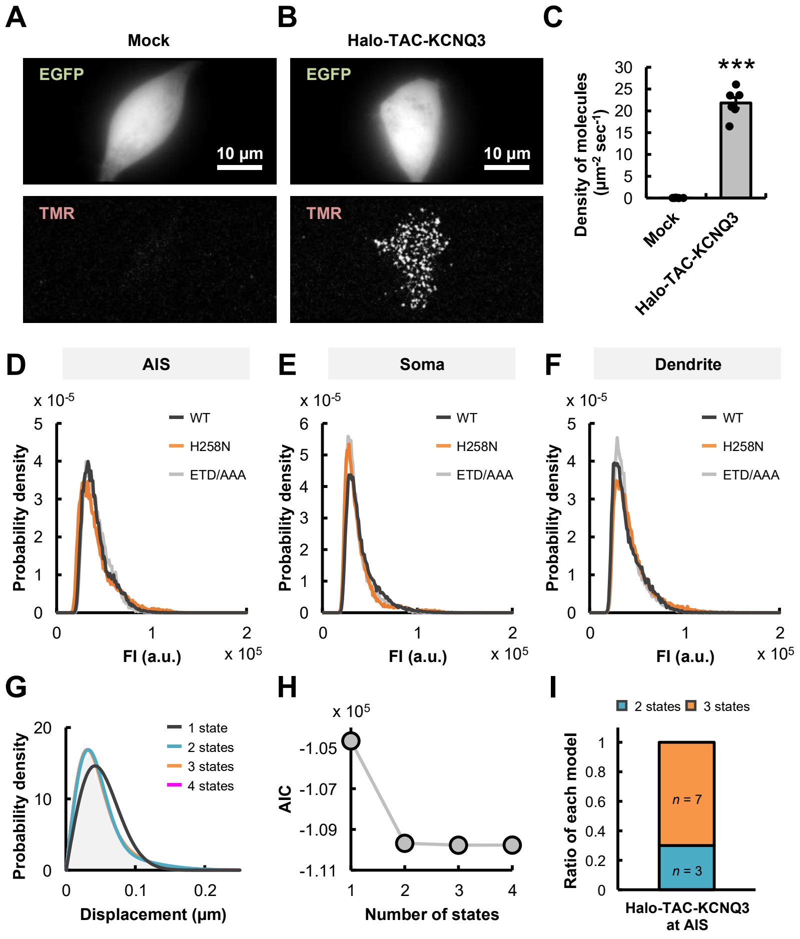
A
Mock
EGFP
10 µm
TMR
B
Halo-TAC-KCNQ3
EGFP
10 µm
TMR
C
***
Density of molecules (µm-2 sec-1)
Mock
Halo-TAC-KCNQ3
D
AIS
E
Soma
F
Dendrite
WT
H258N
ETD/AAA
Probability density
FI (a.u.)
G
1 state
2 states
3 states
4 states
Probability density
Displacement (µm)
H
AIC
Number of states
I
2 states
3 states
Ratio of each model
n = 7
n = 3
Halo-TAC-KCNQ3 at AIS

### Figure S2

**Figure S2: Establishment of single-molecule imaging and the number of diffusion states of Halo-TAC-KCNQ3.**

(A and B) Representative epifluorescence (top, EGFP) and TIRFM (bottom, TMR) images of HEK293T cells expressing EGFP with an empty vector (A) or Halo-TAC-KCNQ3 (B), both subjected to the same TMR labeling procedure. Scale bar, 10 µm. (C) Average density of fluorescent spots per unit time in HEK293T cells expressing an empty vector (Mock, $n$ = 7) or Halo-TAC-KCNQ3 (KCNQ3, $n$ = 6). ***$P$ = 0.000 (Welch's $t$-test). (D-F) Representative probability density distribution of fluorescence intensity of Halo-TAC-KCNQ3 (black, wild-type; orange, H258N; gray, ETD/AAA) at the AIS (D), soma (E), and dendrite (F) in mouse hippocampal neurons. (G) Representative probability density distributions of the displacement ($\Delta t$ = 33 msec) of single Halo-TAC-KCNQ3 molecules at the AIS in neurons. The data were fitted using the 1-, 2-, 3-, and 4-state diffusion models. (H) AIC calculated for each diffusion model in Figure S2G (see "Methods" for Analysis of single-molecule dynamics). In this case, the 3-state model with minimum AIC is estimated as the optimal model. (I) Ratio of neurons where the 2-state model (blue, $n$ = 3) or the 3-state model (orange, $n$ = 7) yielded the minimum AIC values. No neurons were observed where the 1-state model or the 4-state model showed the minimum AIC.

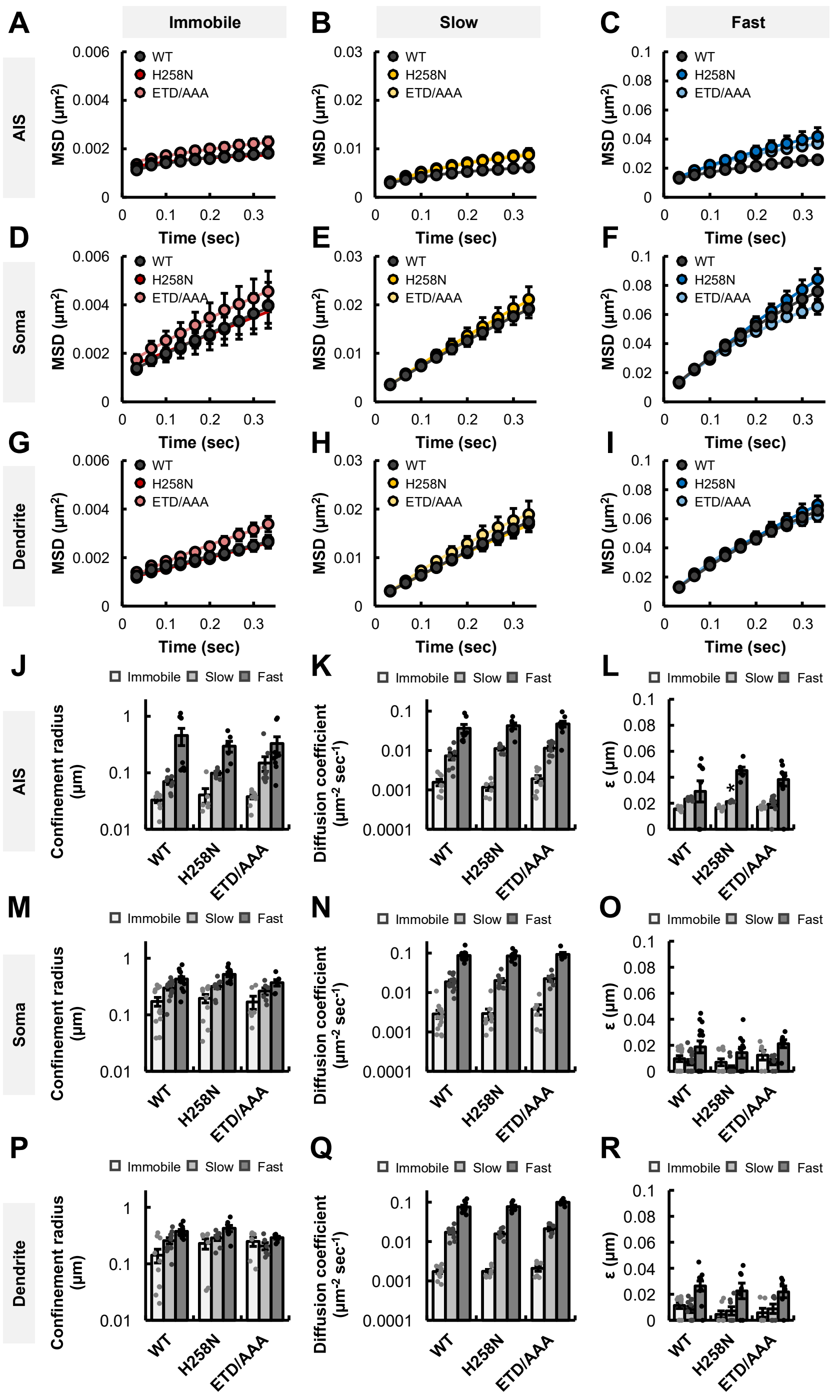

A
Immobile
B
Slow
C
Fast
AIS
Soma
Dendrite
MSD (μm²)
Time (sec)
WT
H258N
ETD/AAA
D
E
F
G
H
I
J
K
L
M
N
O
P
Q
R
Immobile
Slow
Fast
Confinement radius (μm)
Diffusion coefficient (μm² sec⁻¹)
ε (μm)


### Figure S3

**Figure S3: Mean Square Displacement (MSD) analysis of Halo-TAC-KCNQ3 in neurons.** (A-C) Ensemble MSD versus time plots for the immobile (A), slow (B), and fast (C) diffusion state of Halo-TAC-KCNQ3 (wild-type, $n$ = 8; H258N, $n$ = 6; ETD/AAA, $n$ = 9) at the AIS in mouse hippocampal neurons. (D-F) Ensemble MSD versus time plots for the immobile (D), slow (E), and fast (F) diffusing molecules of Halo-TAC-KCNQ3 (wild-type, $n$ = 14; H258N, $n$ = 10; ETD/AAA, $n$ = 6) at the soma in neurons. (G-I) Ensemble MSD versus time plots for the immobile (G), slow (H), and fast (I) diffusing molecules of Halo-TAC-KCNQ3 (wild-type, $n$ = 10; H258N, $n$ = 7; ETD/AAA, $n$ = 6) at the dendrite in mouse hippocampal neurons. (J-L) Confinement radius (J), diffusion coefficient (K), and ε (L) for the immobile (white), slow (light gray), and fast (dark gray) diffusing molecules of Halo-TAC-KCNQ3 (wild-type, $n$ = 8; H258N, $n$ = 6; ETD/AAA, $n$ = 9) at the AIS in neurons, corresponding to Figures S3A-S3C. *$P < 0.05$ (Steel-Dwass test, only comparisons with the "WT" column for the same state are depicted). (M-O) Confinement radius (M), diffusion coefficient (N), and ε (O) for the immobile (white), slow (light gray), and fast (dark gray) diffusing molecules of Halo-TAC-KCNQ3 (wild-type, $n$ = 14; H258N, $n$ = 10; ETD/AAA, $n$ = 6) at the soma in neurons, corresponding to Figures S3D-S3F. $P > 0.05$ for all combinations among wild-type KCNQ3, KCNQ3-H258N, and KCNQ3-ETD/AAA by Steel-Dwass test. (P-R) Confinement radius (P), diffusion coefficient (Q), and ε (R) for the immobile (white), slow (light gray), and fast (dark gray) diffusing molecules of Halo-TAC-KCNQ3 (wild-type, $n$ = 10; H258N, $n$ = 7; ETD/AAA, $n$ = 6) at the dendrite in neurons, corresponding to Figures S3G-S3I. $P > 0.05$ for all combinations among wild-type KCNQ3, KCNQ3-H258N, and KCNQ3-ETD/AAA by Steel-Dwass test. Data are mean ± SE. The data demonstrate that the H258N and ETD/AAA mutations in KCNQ3 have no impact on the confinement radius across all diffusion states in all subcellular regions, and that the diffusion of KCNQ3 is selectively constrained within the AIS regions compared to the somatodendritic regions.

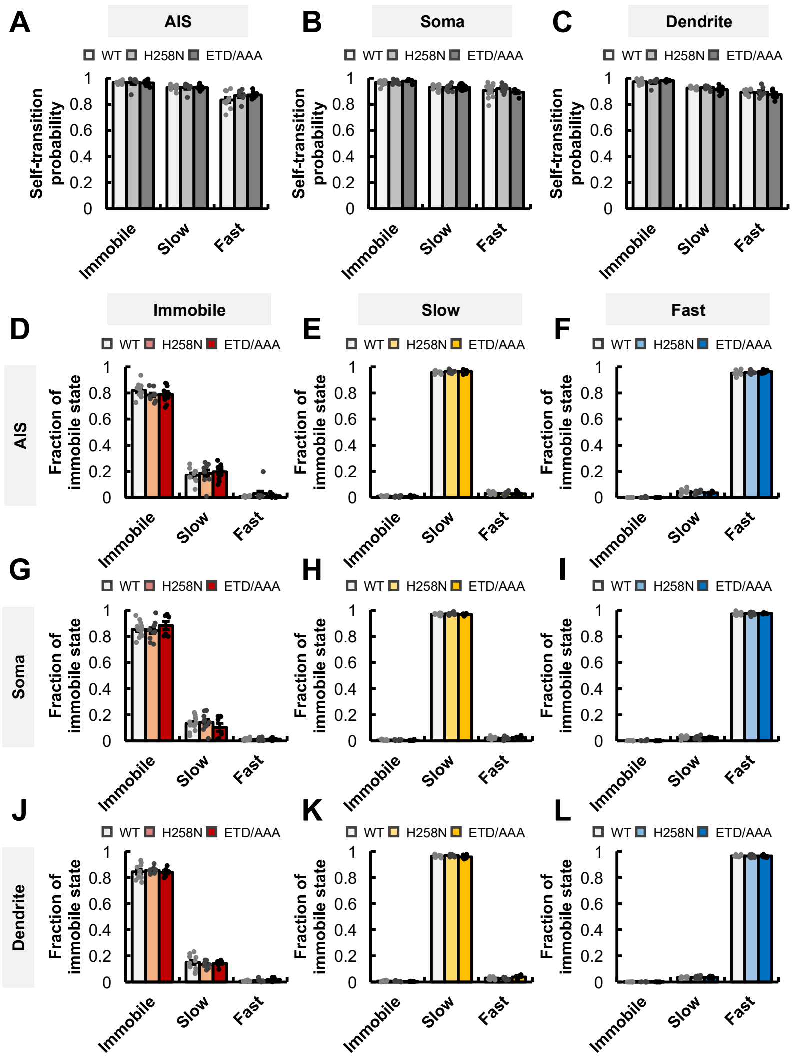
A
AIS
B
Soma
C
Dendrite
WT H258N ETD/AAA
Self-transition probability
1
0.8
0.6
0.4
0.2
0
Immobile
Slow
Fast
D
Immobile
E
Slow
F
Fast
G
H
I
J
K
L
AIS
Soma
Dendrite
Fraction of immobile state

### Figure S4

**Figure S4: Classification of Halo-TAC-KCNQ3 molecules based on the diffusion dynamics.** (A-C) Self-transition probability of immobile, slow-, and fast-diffusion state of Halo-TAC-KCNQ3 at the AIS (A; WT, $n$ = 10; H258N, $n$ = 7; ETD/AAA, $n$ = 9), soma (B; WT, $n$ = 14; H258N, $n$ = 10; ETD/AAA, $n$ = 6), and dendrite (C; WT, $n$ = 10; H258N, $n$ = 7; ETD/AAA, $n$ = 6) in neurons. $P > 0.05$ for all combinations among wild-type KCNQ3, KCNQ3-H258N, and KCNQ3-ETD/AAA by Steel-Dwass test. (D-F) Fraction of each diffusion state in each trajectory of immobile (D), slow (E), and fast (F) diffusing Halo-TAC-KCNQ3 (WT, $n$ = 10; H258N, $n$ = 9; ETD/AAA, $n$ = 9) molecules at the AIS in neurons. $P > 0.05$ for all combinations among wild-type KCNQ3, KCNQ3-H258N, and KCNQ3-ETD/AAA by Steel-Dwass test. (G-I) Fraction of each diffusion state in each trajectory of immobile (G), slow (H), and fast (I) diffusing Halo-TAC-KCNQ3 (WT, $n$ = 14; H258N, $n$ = 10; ETD/AAA, $n$ = 6) molecules at the soma in neurons. $P > 0.05$ for all combinations among wild-type KCNQ3, KCNQ3-H258N, and KCNQ3-ETD/AAA by Steel-Dwass test. (J-L) Fraction of each diffusion state in each trajectory of immobile (J), slow (K), and fast (L) diffusing Halo-TAC-KCNQ3 (WT, $n$ = 10; H258N, $n$ = 7; ETD/AAA, $n$ = 6) molecules at the dendrite in neurons. $P > 0.05$ for all combinations among wild-type KCNQ3, KCNQ3-H258N, and KCNQ3-ETD/AAA by Steel-Dwass test. Data are mean ± SE.

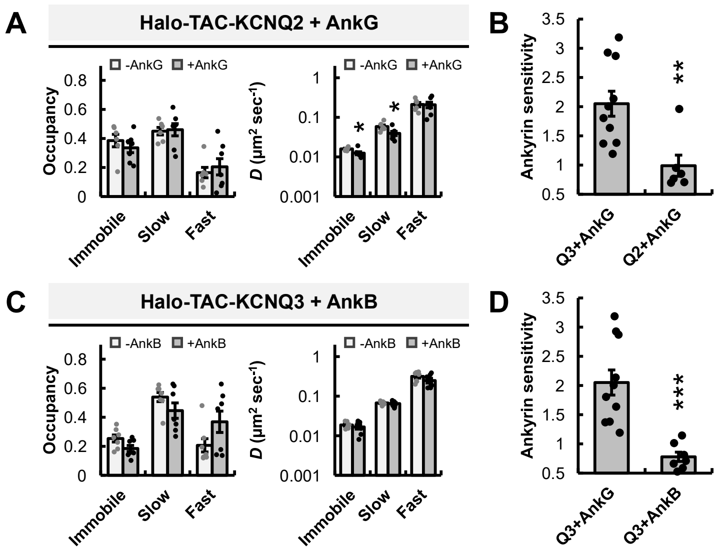
A
Halo-TAC-KCNQ2 + AnkG
-AnkG
+AnkG
Occupancy
0.8
0.6
0.4
0.2
0
Immobile
Slow
Fast
D (μm² sec⁻¹)
1
0.1
0.01
0.001
B
Ankyrin sensitivity
3.5
3
2.5
2
1.5
1
0.5
**
Q3+AnkG
Q2+AnkG
C
Halo-TAC-KCNQ3 + AnkB
-AnkB
+AnkB
D
***
Q3+AnkG
Q3+AnkB

### Figure S5

**Figure S5. Specificity of interaction between KCNQ3 and ankyrin-G.**
(A) Occupancy (left) and diffusion coefficient (right) for immobile, slow-, and fast-diffusion state of Halo-TAC-KCNQ2 in HEK293T cells with (+AnkG, $n$ = 7) and without (-AnkG, $n$ = 6) 270 kDa ankyrin-G. *$P$ < 0.05 (Welch's $t$-test, comparisons were made with the "-AnkG" group for the same state). (B) Ankyrin sensitivity of Halo-TAC-KCNQ2 and Halo-TAC-KCNQ3 calculated as the ratio between the immobile state occupancy with and without 270 kDa ankyrin-G (Q3+AnkG, $n$ = 10; Q2+AnkG, $n$ = 6). **$P$ = 0.004 (Welch's $t$-test). (C) Occupancy (left) and diffusion coefficient (right) for immobile, slow-, and fast-diffusion state of Halo-TAC-KCNQ3 in HEK293T cells with (+AnkB, $n$ = 7) and without (-AnkB, $n$ = 7) 220 kDa ankyrin-B. No significant differences were detected in all parameters with and without ankyrin-B (Welch's $t$-test, comparisons were made with the "-AnkB" group for the same state). (D) Ankyrin sensitivity of Halo-TAC-KCNQ3 calculated as the ratio between the immobile state occupancy with and without 270 kDa ankyrin-G or 220 kDa ankyrin-B (Q3+AnkG, $n$ = 10; Q3+AnkB, $n$ = 7). ***$P$ = 0.000 (Welch's $t$-test). Data are mean ± SE. These data suggest that KCNQ3 has a higher affinity for ankyrin-G in comparison to KCNQ2. Furthermore, it is indicated that KCNQ3 exhibits a greater sensitivity to ankyrin-G rather than ankyrin-B.

#### Supplementary Table

**Table S1. Fraction of immobile, slow-, and fast-diffusing Halo-TAC-KCNQ3.**

The number of immobile, slow-, and fast-diffusing Halo-TAC-KCNQ3 molecules appearing per 10 $\mu m^2$ over 15 sec was summed, and the fraction of each classified group was calculated (WT at the AIS, 10 cells; H258N at the AIS, 9 cells; ETD/AAA at the AIS, 9 cells; WT at the soma, 14 cells; WT at the dendrite, 10 cells). *P*-values were obtained from Chi-square test and subsequent residual analysis.

**Table S1: Fraction of immobile, slow-, and fast-diffusing molecules**

| | Fraction of molecules (number of molecules) | | |
|---|---|---|---|
| Data | Immobile | Slow | Fast |
| WT at AIS | 0.063<br>(93) | 0.668<br>(985) | 0.269<br>(396) |
| H258N at AIS | 0.028<br>(20) | 0.630<br>(449) | 0.342<br>(244) |
| ETD/AAA at AIS | 0.032<br>(22) | 0.636<br>(444) | 0.332<br>(232) |

| | Comparison | | |
|---|---|---|---|
| | WT-H258N | WT-ETD/AAA | H258N-ETD/AAA |
| *P*-value (Chi-square test) | 0.000 | 0.000 | 0.875 |

| | *P*-value (Residual analysis after Chi-square test) | | |
|---|---|---|---|
| Comparison | Immobile | Slow | Fast |
| WT-H258N | 0.001 | 0.076 | 0.000 |
| WT-ETD/AAA | 0.002 | 0.140 | 0.002 |
| H258N-ETD/AAA | 0.702 | 0.804 | 0.696 |

| | Fraction of molecules (number of molecules) | | |
|---|---|---|---|
| Data | Immobile | Slow | Fast |
| WT at AIS | 0.063<br>(93) | 0.668<br>(985) | 0.269<br>(396) |
| WT at Soma | 0.038<br>(25) | 0.573<br>(378) | 0.389<br>(257) |
| WT at Dendrite | 0.034<br>(19) | 0.613<br>(343) | 0.354<br>(198) |

| | Comparison | | |
|---|---|---|---|
| | WT-H258N | WT-ETD/AAA | H258N-ETD/AAA |
| *P*-value (Chi-square test) | 0.000 | 0.000 | 0.371 |

| | *P*-value (Residual analysis after Chi-square test) | | |
|---|---|---|---|
| Comparison | Immobile | Slow | Fast |
| AIS-Soma | 0.018 | 0.000 | 0.000 |
| AIS-Dendrite | 0.010 | 0.018 | 0.000 |
| Soma-Dendrite | 0.712 | 0.159 | 0.197 |

#### Description of Supplementary Videos

**Video S1. Single-molecules of TMR-labeled Halo-TAC-KCNQ3 at the AIS in neurons.**
Time-lapse TIRFM images depict individual molecules of TMR-labeled Halo-TAC-KCNQ3 (left, WT; center, H258N; right, ETD/AAA) on the cell surface at the AIS. These images correspond to Figures 5 and 6. The images are displayed at intervals of 33 msec. Time, sec. Scale bar, 1 µm.

**Video S2. Single-molecules of TMR-labeled Halo-TAC-KCNQ3 at the soma in neurons.**
Time-lapse TIRFM images depict individual molecules of TMR-labeled Halo-TAC-KCNQ3 (left, WT; center, H258N; right, ETD/AAA) on the cell surface at the soma. These images correspond to Figures 5 and 6. The images are displayed at intervals of 33 msec. Time, sec. Scale bar, 1 µm.

**Video S3. Single-molecules of TMR-labeled Halo-TAC-KCNQ3 at the dendrite in neurons.**
Time-lapse TIRFM images depict individual molecules of TMR-labeled Halo-TAC-KCNQ3 (left, WT; center, H258N; right, ETD/AAA) on the cell surface at the dendrite. These images correspond to Figures 5 and 6. The images are displayed at intervals of 33 msec. Time, sec. Scale bar, 1 µm.

**Video S4. Single-molecules of TMR-labeled Halo-TAC-KCNQ3 in HEK293T cells.**
Time-lapse TIRFM images depict individual molecules of TMR-labeled Halo-TAC-KCNQ3 (left, without ankyrin; center, with 270 kDa ankyrin-G; right, with 220 kDa ankyrin-B) on the surface of HEK293T cells. These images correspond to Figures 7 and S5. The images are displayed at intervals of 33 msec. Time, sec. Scale bar, 5 µm.

**Video S5. Single-molecules of TMR-labeled Halo-TAC-KCNQ2 in HEK293T cells.**
Time-lapse TIRFM images depict individual molecules of TMR-labeled Halo-TAC-KCNQ2 (left, without 270 kDa ankyrin-G; right, with 270 kDa ankyrin-G) on the surface of HEK293T cells. These images correspond to Figure S5. The images are displayed at intervals of 33 msec. Time, sec. Scale bar, 5 µm.

**Video S6. Single-molecules of TMR-labeled Halo-TAC-KCNQ3 mutants in HEK293T cells.**
Time-lapse TIRFM images depict individual molecules of Halo-TAC-KCNQ3-ETD/AAA (upper left), Halo-TAC-KCNQ3-FDQ/RNE (upper right), Halo-TAC-KCNQ3-Y267C (lower left), and Halo-TAC-KCNQ3-H258N (lower right) with and without 270 kDa ankyrin-G on the surface of

HEK293T cells. All Halo-TAC-KCNQ3 mutants were labeled with TMR. These images correspond to Figure 7. The images are displayed at intervals of 33 msec. Time, sec. Scale bar, 5 µm.
